# Curcumin Conjugated Carbon Quantum Dots: A Theranostic Probe to Study BSA Interaction, Cellular Imaging, and RICS-Based Intracellular Transport

**DOI:** 10.1101/2025.06.24.661223

**Authors:** Arun K. Upadhyaya, Dibyendu K. Sasmal

**Affiliations:** Department of Chemistry, Indian Institute of Technology Jodhpur, India

**Keywords:** Carbon Quantum Dots, Protein Corona, Fluorescence Quenching, Raster Image Correlation Spectroscopy (RICS), antibacterial activity, Cytotoxicity, Cellular Uptake

## Abstract

Plant-based products such as jamun leaves and herbaceous perennials like turmeric are replete with carbon-rich elements, and our recent focus has been directed to synthesizing Carbon Quantum Dots (CQDs) in a low-cost, straightforward, and sustainable method, with their biological and therapeutic applications. A valuable insight we have gained from our results is the environmentally friendly production of CQDs with their distinct physicochemical characteristics, in addition to excellent water solubility and remarkable stability. The as-synthesized CQDs manifest good quantum yields with many potential applications. Moreover, for the stability and enhancement of the solubilization of the hydrophobic drug curcumin, to improve its uptake and bio accessibility, CQDs are a unique choice. After the conjugation of CQDs with curcumin (C-CQDs), the findings we have obtained with partially unfolded BSA demonstrate a modest interaction involving dynamic quenching, as exemplified by steady-state and time-resolved fluorescence quenching driven by Forster Resonance Energy Transfer (FRET). In addition to that, our research also delved into the application of *in vivo* bioimaging, and by using that fluorescence imaging technique, we introduced Raster Image Correlation Spectroscopy (RICS) techniques to find the diffusion of both CQDs and C-CQDs in the live cell. The study also reaffirms the inherent biocompatibility and antibacterial ability confirmed by the MTT-based cytotoxicity assay and MIC study.

## Introduction

In recent years, due to the remarkable photoluminescence, low toxicity, aqueous solubility, and surface tunability, Carbon Quantum Dots (CQDs) have gained considerable attention in the field of nanobiotechnology.^1–2^ Among the various nanomaterials explored, these zero-dimensional carbon-based nanomaterials offer significant advantages over traditional semiconductor quantum dots, especially in biological applications, where biocompatibility and environmental safety are crucial.^3–4^ Factors, such as fluorescence properties, functional groups, and sizes of less than 10 nm, empower CQDs to carry various hydrophobic drugs. Along with the versatile surface chemistry, it allows them for easy conjugation with therapeutic molecules, making them well-suited for drug transport and bioimaging uses. The ultrasmall size of CQDs (below 10 nm) can easily cross the biological barrier, which helps to deliver the drugs much more effectively to the cells.^5^ Nowadays, researchers have used CQDs extensively in cancer cell imaging and drug delivery.^6–7^ Hydrophobic drugs like Doxorubicin (DOX), Gemcitabine (Gem), Curcumin, paclitaxel, hyaluronan, and cisplatin can be easily bound to CQD’s surface because of their unique surface chemistry. As per a report by Yang *et al.*, the CQD-DOX complexes induce apoptosis in human lung adenocarcinoma cells.^8^ Comparably, Winer *et al.* found that CQD-bound Gemcitabine is mainly effective against pancreatic cancers, especially those undergoing tumor resections.^9^ A study by Ajima *et al.* reported that the conjugation of cisplatin with carbon nanotubes (CNTs) had anti-cancer efficiency by 6 times.^10^

Apart from various hydrophobic drugs, curcumin (1,7-bis(4-hydroxy-3-methoxyphenyl)-1,6-heptane-3,5-dione), a natural polyphenol found in turmeric, is used for the treatment of antimicrobial, anti-inflammatory, antioxidant, and anticancer properties as well.^11^ However, because of its low solubility in water, it limits its absorption and solubility inside the human body under basic pH and neutral conditions, even with a promising therapeutic effect.^12^ From several studies, it has been found that Curcumin can suppress the growth of cancer cells in the liver, stomach, kidney, breast, and brain as well.^13^ A type of protein kinase known as cyclin-dependent kinases (CDKs), which control the progression of the cell cycle via phosphorylation of enzymatic and structural targets, can be reduced by Curcumin.^14^ Despite having some excellent medical development, one of the major drawbacks of curcumin is its low bioavailability, physiological pH instability, lower solubility, quick consumption, short biological half-life, and fast systemic removal, which make curcumin a less or undetectable amount in extra-intestinal tissues and blood. Also, from animal models, it has been found that around 90% of the curcumin is primarily removed if given orally.^15^

However, to resolve those challenges made by curcumin, nanotechnology plays a role for improving the nano formulation techniques, somehow increasing the bioavailability and solubility.^16^ Nanomaterials like Liposomes and graphene oxide have been used as effective carriers of the drug Curcumin.^17–18^ In a similar way, MSN (Mesoporous silica Nanoparticle) also helps to bind Curcumin with GaNO_3_ and fucoidan.^19–20^ Cyclodextrin nanoparticles help to carry curcumin to the cancerous cells.^21^ Peptides are also carrying the drug Curcumin with DOX to the cancerous cells.^22^ Biodegradability and biocompatibility can also be increased by some of the biopolymers like starch, alginate, chitosan, and silk, which can also help curcumin to carry up to the cancer cells.^23^

In addition to that, due to the specific physical and chemical properties of the quantum dots, several studies have been conducted to understand and explore how the quantum dots affect the essential macromolecules like serum albumin protein (BSA).^24–25^ With a molecular weight of 66.5 kDa, globular protein BSA comprises 583 amino acids, 17 cysteine residues connected by 8 disulfide bridges with one free thiol group.^26^ Due to their ability to bind reversibly with therapeutic drug molecules, they play a vital role in drug delivery and drug carriers.^27^ With almost 80% sequence homology, BSA is also structurally homologous to human serum albumin (HSA).^28^ Among various serum albumins, BSA is widely selected as a model protein for extensive study because of its low cost, easy availability, and analogous nature to HSA.^29^ However, monitoring the BSA fluorescence quenching with the addition of quantum dots uncovers the interactions between them. A scarcity of exploration from the literature is the interaction between BSA and quantum dots, promising numerous applications in biological research. A study by Yin *et al.* shows the interaction of serum albumin protein with water-soluble ultrasmall gold nanoparticles, where, from a spectroscopic approach, a hard protein corona formation was confirmed.^30^ A recent study by Agarwala *et al* reported the hydrophobic interaction of bovine serum albumin (BSA) with curcumin by using the spectroscopic techniques.^31^

An extended version of FCS, called Raster Image correlation spectroscopy (RICS), which uses raster scanning to capture the fluctuations in the intensity caused by the movement of fluorescent molecules by using a confocal laser scanning microscopy (CLSM).^32^ Specifically, this method is particularly advantageous for studying heterogeneous environments where traditional FCS may fail to capture spatial variations. Through the detection volume, diffusion of fluorescent particles can be measured by measuring one pixel’s intensity for a very particular time period, by measuring the adjacent pixel’s intensity immediately, and then, within each frame, it can be correlated pairwise to dynamically characterise the decay processes.^33^ However, the spatial correlation can depend on the size of the pixel, pixel dwell time, and the rate of diffusion as well.^33^ RICS has been applied in diverse fields such as biophysics, soft matter physics, and nanomedicine, providing insights into membrane protein interactions, intracellular transport mechanisms, and drug delivery systems.^34–35^

In a broader contrast to this work, the study manifests the synthesis of fluorescent, water-soluble Carbon Quantum Dots (CQDs) using the reflux method in an eco-friendly way. Using many spectroscopic methods, we have characterized the CQDs, and significantly, we have obtained exceptionally stable quantum dots having a quantum yield of 36%. Additionally, we have synthesized curcumin-conjugated CQDs (C-CQDs), which show excellent water solubility. To understand the interaction between Serum albumin protein, BSA, and formed C-CQDs, we have studied the fluorescence quenching experiment, where we have found that the proteins weakly adhere in their hydrophobic sites to the surface of CQDs, where curcumin is there. Apart from that, both CQDs and C-CQDs, because of their fluorescence properties and biocompatible nature, can be used to visualize the human lung cancer cell (A549) as a bioimaging probe. We have also taken the imaging-based diffusion study, Raster Image Correlation Spectroscopy (RICS), to find out the diffusivity of both the fluorescent probe CQDs and C-CQDs in the live cancer cell A549. Additionally, using MTT-based cytotoxicity assay and MIC assay, we have studied the cell viability and antibacterial properties, respectively.

### Experimental Section

#### Chemicals and Reagents

For the synthesis of Carbon Quantum Dots (CQDs), Jamun leaves (Syzygium cumin) have been collected from inside the IIT Jodhpur campus, and raw turmeric (Curcuma longa) has been purchased from the local market near Jodhpur. Then, by using reflux methods, CQDs have been synthesized. From Sigma-Aldrich (USA), Bovine Serum Albumin (BSA) and Curcumin were purchased. From Loba Chemie Pvt. Ltd. (India), Phosphate-buffered saline (PBS) tablets (pH 7.4) were purchased. High-purity tris-hydroxymethylaminomethane (Tris-HCl) was purchased from Fisher Scientific (India). Dulbecco’s modified Eagle medium (DMEM) and Trypsin-EDTA solution were purchased from Himedia. From Sigma-Aldrich (USA), Penicillin-Streptomycin and Thiazolyl Blue Tetrazolium Bromide (MTT) were purchased. Ethanol was bought from Grace LifeTech Pvt. Ltd. No further purification of the chemicals was needed, as they are all analytical grades. Throughout the analysis, MILLI-Q water was used.

#### Green Synthesis of CQDs

Fresh green Leaves of Syzygium cumini and about 100 gms of raw Curcuma longa were taken and first washed properly with normal water, followed by DI water. It was then cut into small pieces and crushed finely. The juice from the raw materials was extracted and filtered by using a Whatman filter paper to get a clear solution, and from that clear solution, 15 mL of leave juice and 5 mL of Curcuma longa juice were taken in a 50 mL round bottom neck flask (3:1 mixture) and refluxed for 3.5 hours at 210° C, resulting in a brown solution. The residue was cooled at room temperature, and then around 30 mL of Dichloromethane (DCM) was added to that residue to remove the unreacted organic molecules. Afterward, by using Whatman filter paper, the upper aqueous layer was filtered. It was then centrifuged at 14,500 rpm for 30 minutes to remove the larger particles as precipitate. CQDs with smaller particle sizes were then dried at 60° C by placing them on top of a hot plate in a dark place, followed by a vacuum desiccator to get the powder form of CQDs. It was then stored in a glass bottle covered with aluminium foil in a dark environment for further characterization and applications.

#### Synthesis of Curcumin-loaded CQDs (C-CQDs)

Solution of CQDs was made with 2 μg/mL of solid CQDs in DI water. From that solution, 2 mL was taken and added to 1 mg of curcumin. The mixture was stirred for 3 hours in the dark using a magnetic stirrer. To pelletize the unbound curcumin, for 15 minutes at 1200 rpm, it was then centrifuged. The unbound curcumin was separated by centrifugation, and a precipitate was formed. Further, at 14500 rpm, it was centrifuged to get the Curcumin bound with CQDs (C-CQDs). It was then separated and placed inside a hot oven at 60° C and then under a vacuum overnight to remove moisture and then stored in the dark. The C-CQDs pellet was dispersed in MILLI-Q water whenever needed in a concentration of 1 mg/mL. Then, for further characterization and applications, we synthesized that on a larger scale.

#### UV-Visible Absorption Spectroscopy and Fluorescence Spectroscopy

UV-visible absorption spectra and fluorescence emission spectra were obtained by using a JASCO Spectrophotometer IX800 and a JASCO Spectrofluorometer FP-8300, respectively, using different quartz cuvettes of the same path length of 1 cm to determine the absorption and emission properties of the samples. The scan speed for each spectrum was kept at 1000 nm/min. For emission spectra, the response time was kept at 50 ms, and the excitation and emission bandwidths were set at 5 nm. As a reference standard, MILLI-Q water was used for each set of experiments with CQDs and C-CQDs. For the excitation and emission spectra of BSA only, Tris buffer was used as a reference. To study the excitation-dependent emission spectrum, different ranges of excitation spectra have been taken to collect the emission spectra. For the calculation of concentrations of BSA, equation 1 was used.^36^

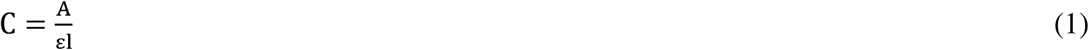

where A. Absorbance, ε . Molar Absorptivity (For BSA, we used ε = 43824Lmol^-1^cm^-1^ at excitation 280 nm)36, l . the path length of the cuvette (1 cm).

#### 3D Fluorescence Spectroscopy

With a scan speed of 1000 nm/min and response time of 50 ms at low sensitivity, 3D fluorescence spectra were recorded by taking 5 μL of stock CQDs and C-CQDs in 2 mL of water, separately using a JASCO Spectrofluorometer FP-8300 at excitation spectra of 300 nm to 600 nm, both for CQDs and C-CQDs and the emission range collected for CQDs and C-CQDs was 310-700 nm. The excitation and emission bandwidths are set as 10 nm for each sample.

#### FTIR spectroscopy

Fourier-transform infrared (FTIR) spectra were recorded using a Bruker Alpha II FTIR spectrometer with an Attenuated total reflectance (ATR) accessory over the range of 600-4000 cm^-1^ by taking 100 µl of each sample for the determination of various surface functional groups. For blank measurement, only MILLI-Q water was taken.

#### Raman Spectroscopy

A solid, dry sample of CQDs and C-CQDs was taken on a glass slide, with a 532 nm laser, and by using a Bay Spec Raman Spectrophotometer, Raman Spectra were collected. Also, in the case of curcumin, a pinch of solid curcumin was taken on the surface of a glass slide for the spectra. For the collection of Raman Spectra, the Raman shift for each of the samples was taken from 300-2000 cm^-1^.

### FESEM Analysis

To know the morphology and microstructural properties, a Field Emission Scanning Electron Microscopy (FESEM) image was collected using Thermo Fisher Scientific Apreo 2S Scanning Electron Microscopy. For the sample preparation, dry and solid CQD was taken on the surface of a carbon tape. The image was taken for the analysis, keeping the magnification as 5,0,000X and HV as 15 kV.

#### Energy Dispersive X-ray (EDX) Analysis

To know the elemental mapping and Energy dispersive X-ray analysis, a Carl Zeiss-EVO 18 Scanning Electron Microscope (SEM) with EDS-51-ADDD-0048 was used. For the study, solid and dry CQDs were placed on carbon tape and taken for analysis. On the next day, after the sample was completely dry, the samples were taken for the study.

#### Dynamic Light Scattering (DLS) Analysis

To know the hydrodynamic size, zeta potential, and electrophoretic mobility, DLS measurement was done using Zetasizer Pro ZSU3200 in an aqueous medium. For the calculation of size, 5 μL of CQDs was added to 2 mL of DI water as a dispersant and was taken inside a disposable 10 × 10 plastic cell (DTS0012) for the measurement, keeping the temperature as 25°C with an equilibration time of 120 sec. Similarly, for zeta potential and electrophoretic mobility, the same sample was taken inside a disposable folded capillary cell (DTS1080) at the same temperature and equilibration time. For the C-CQD, to know the hydrodynamic size, 5 mL of water dispersed sample was taken inside a disposable 10 ×10 plastic cells (DTS0012).

#### XRD Analysis

Powdered XRD can be analyzed by using a Panalytical Empyrean high-resolution X-ray diffractometer, keeping 2θ values ranging from 5 to 60 degrees. The as-prepared carbon quantum dots in their powder form of about 100 mg were kept on the surface of the sample chamber, and the diffraction pattern was taken.

#### X-Ray Photoelectron Spectroscopy (XPS) Analysis

To know the elemental bonding and chemical states of the prepared sample (CQDs and C-CQDs), X-ray photoelectron spectroscopy (XPS) (PHI-5000 VersaProbe III; UlvacPhi, Inc.) was used. Under ultra-high vacuum [2 × 10^-7^] Pa, a monochromatic AlKa anode source with high energy (1.486 keV) was used to generate X-rays. The step width was set as 0.05 eV, and the pass energy for the survey spectrum was kept at 187.5 eV, whereas for the core level, to get better resolution, at a low set of pass energies of 23.75 eV, the spectrum was acquired. For peak shift correction of the constituent elements, the C1s reference used in this spectrum is 284.5 eV. During data acquisition, an electron neutralizer (E-Neut) was used for charge neutralization on the sample surface. To obtain a better fit, both Gauss and Lorentz types of functions were used in the curve fitting procedure. With isopropanol, the holder of the sample was adequately cleaned before measuring.

#### Steady-State and Time-Resolved Fluorescence Measurement

To investigate the interaction between BSA and C-CQD, we conducted steady-state fluorescence spectroscopy and time-correlated fluorescence measurements. First, we prepared a stock solution of BSA with a molecular weight of 66 kDa by dissolving an appropriate amount of BSA in 50 mM Tris buffer at pH =7.4. Then, the solution was stored at 4 ^0^C for further experiments. For the Steady-state quenching experiments, we prepare 25 μM concentration of BSA from the stock, and in that solution, the C-CQD was added from 0.5 μL to 22 μL gradually, and the fluorescence was collected using JASCO Spectrofluorometer FP-8300. All the measurement was performed at an excitation wavelength of 280 nm, and the emission was collected from 290 nm to 600 nm. For the calculation of the binding constant or association constant for enhancement of overall fluorescence upon the addition of the quencher, we applied the Benesi-Hildebrand equation :^37^

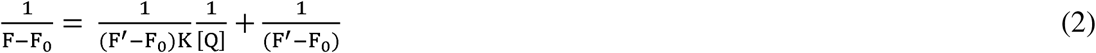

of Q (quencher), *F*’ = Fluorescence intensity of the fluorophore at the highest concentration of Where F and F_0_ = Steady-state fluorescence intensity of fluorophore in the presence and absence quencher, [Q] = Concentration of the quencher (C-CQD), K = Binding constant, which was obtained by the equation:

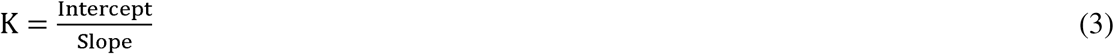

To calculate the Stern-Volmer quenching constant and stoichiometry of interactions, we used Stern-Volmer and double logarithm Stern-Volmer (DLSV), respectively given by equations (4) and (5).

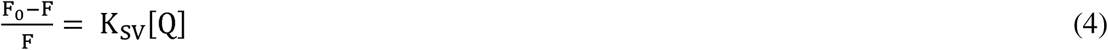

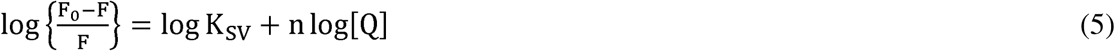

where, K_sv_ = Stern-Volmer quenching constant and n = number of binding sites. The values K_sv_ and n were obtained from the slopes of the plot of 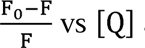 and 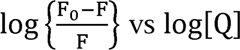. Further, we have explored the luminescence lifetime of 25 μM BSA in the presence and absence of C-CQDs of different concentrations in Tris buffer (pH=7.4). By using a picosecond diode, the experiment was conducted using the Edinburgh FLS 1000, with an LED source at excitation of 280 nm, a full width half maximum value of 900 ps, and a spectral width of 10 nm. Then, the experimentally obtained data are fitted using a bi-exponential function represented by the equation below.

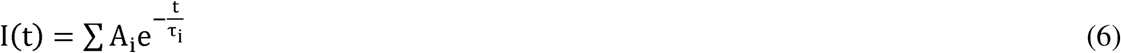

where A_i_ represents the amplitude of the i^th^ lifetime associated with the i^th^ lifetime τ_i_. The average lifetime <τ> was calculated by the equation.

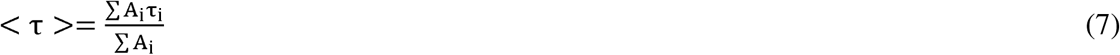

#### Forster Resonance Energy Transfer (FRET)

With respect to the acceptor rise, the Fo rster energy transfer efficiency from donor to acceptor was calculated by the equation:^36^

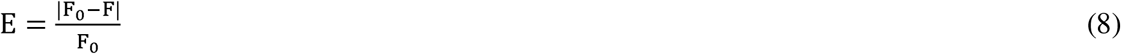

Where F_0_ and F respectively represent the intensities of fluorescence for the acceptor in the absence and presence of the donor. To get a positive increase in acceptor fluorescence intensity, the modulus of (F_0_ – F) was taken.

#### Adsorption Efficiency Calculation

The adsorption efficiency was calculated using the following equation^38^

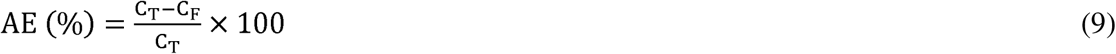

Where C_T_ is the total added curcumin and C_F_ is the curcumin present after centrifugation at 1200 rpm, which is not bound.

#### Raster Image Correlation Spectroscopy (RICS) Analysis

To get Raster scan images, accord with the pixel size of 50 nm, images were collected with a aspect ratio of 256 × 256 pixels using a 100X objective at 9.9X zoom and by FluoView imaging software (Olympus FluoView 1000 Laser Scanning Confocal Microscope). A 405-nm laser was used for the excitation of samples, and the pixel dwell time was set as 12.5 μs/pixel to scan inside the cell. To subtract the background and improve the signal-to-noise ratio, multiple stacks of images were captured for each set of experiments with no delay between the frames. Here, we have taken 100 stacks of images for each set of experiments. For the imaging experiment, 10 μL of CQDs and C-CQDs were added separately to 1 mL of Imaging media as discussed in the imaging part. Initially, the image was taken in an aspect ratio of 512 × 512, and then for the RICS experiment, with 9.9X zoom, a stack of 256×256-pixel images was taken.

#### Diffusion Coefficient Measurement using RICS

For the calibration of beam waist (ω_0_), first, we have taken 50 nM of Rhodamine-6G with a known diffusion coefficient of 2.8 × 10^-6^ cm^2^/sec in water.^39^ Then, using SimFCS 4 software and with the known parameters of pixel dwell time, line time, retracing time, size of each pixel, and the beam waist, the diffusion coefficient was measured. For each stack, autocorrelation was calculated as an average of all the images using an equation.^33^

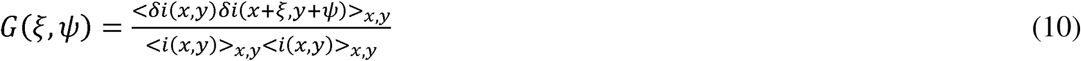

From the above equation, ξ and ψ represents the x and y spatial correlation shifts, *i(x, y)* is pixel intensity of the image, and .δ*i*= *i* –〈…〉, and 〈…〉_x,y_. After averaging each frame’s image correlation. *G_RICS_*(ξ, ψ) is separated into two parts: like *G*(ξ, ψ) and *S*(ξ, ψ), where correlations, the result was fitted with an equation that links the diffusion coefficient *G*(ξ, ψ) related to molecular dynamics and *S*(ξ, ψ) related to scanning optics. The most basic form of the autocorrelation function for 3D diffusion is:

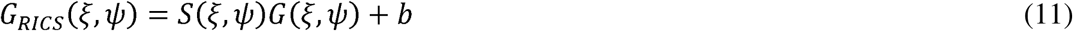

Where,

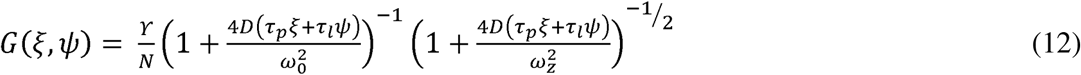

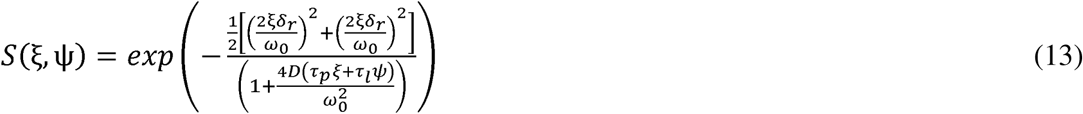

With D as a diffusion coefficient and N as the particle numbers, τ*_p_* represents the pixel dwell time with the line time as τ*_l_* and the pixel size as τ*_r_*. From the equation, ω*_0_* represents the beam waist, and b is the background.

#### Cell Culture and Imaging Experiment

Human lung adenocarcinoma epithelial cell A549 was cultured in Dulbecco’s modified Eagle medium containing 4.5 gm/L glucose (Himedia, India), with 10% fetal bovine serum (Himedia, humidified incubator with a 5% carbon dioxide environment, the cells were grown at 37°C India) and 100-unit penicillin-streptomycin (Sigma Aldrich) in a tissue culture flask. In a temperature. For the microscopy experiment, a 22 ×22 µm clean coverslip was placed on a 35 mm ×12 mm petri dish, and the cells were seeded for 24 hours in a phenol red-free culture media. To observe the effect of CQDs and C-CQDs on the cell imaging, 10 μ*L* of CQDs and C-CQDs was added to different petri dishes, respectively, and then 1 mL of culture media was added. For the time-dependent cellular uptake study, five different petri dishes with cells were taken, and the same amount (10 μ*L*) of C-CQDs was added to each of the petri dishes. The cells were then incubated with C-CQDs at different time scales. Then, the strained cell was washed with PBS buffer to remove the excess C-CQDs. The coverslip was carefully removed and mounted over a cleaned glass slide for microscopy imaging. With a 100X objective (Numerical aperture 1.45), using an Olympus FV1000 Laser Scanning Confocal Microscope for focusing the excitation light onto the cover glass, all the bioimaging studies were done.

#### Preparation of the Cell Sample for FESEM

We have taken three different types of Petri dishes, where A549 cells were seeded first for about 3-4 days to become 70-80% confluent on the surface of the glass slide. Among these three, one petri dish was taken for the control, where no CQDs or C-CQDs had been added. In the other two petri dishes, we have added 2 μ*L* of the CQDs and C-CQDs to each of the petri dishes. The cells were then cultured for 30 minutes with both CQDs and C-CQDs in phenol red-free DMEM media to permit the cells to enter the cells. Then, using 4% paraformaldehyde solution, the cells were fixed, followed by washing three to four times with PBS buffer. Then, with 10-minute intervals, the coverslips were dehydrated by using 50%, 70%, 90%, and 100% ethanol for 5 minutes each. The coverslips were then stored inside the desiccator.

#### EMCCD Cell Imaging

For two-channel imaging of Quantum Dots (CQDs and C-CQDs) and DAPI, we used a halogen lamp for excitation and an Andor EMCCD camera for imaging. Imaging for CQDs and C-CQDs uptake, which is falling in the excitation and emission range of Green fluorescent protein (GFP), was collected by using the GFP channel from the halogen lamp in response to 420 nm excitation and > 500 nm emission. For the GFP channel, the camera exposure time was set constant at 100 ms with an EM-Gain of 50 for all sets of experiments. The fluorescence emission from DAPI in the range of 440–500 nm was collected in response to 358 nm excitation by a halogen lamp. For all sets of experiments, the camera exposure time was set at 20 ms with EM Gain at 20 for the DAPI channel.

#### MIC study of C-CQDs

Using liquid LB medium, the bacterial strain (Escherichia coli DH5α) was incubated at 37 °C overnight with continuous mechanical stirring at 120 rpm. On the next day, the tube containing the bacteria with media is centrifuged for 3 minutes at 6000 rpm to get the bacterial pellet. The pellet was then mixed with 2 mL of fresh LB media to make it homogeneous. The media with bacteria was then equally distributed in different tubes containing fresh media. It was allowed to grow, and then various concentrations of CQDs and C-CQDs were added. All these tubes were incubated overnight at 37 °C, 120 rpm with continuous stirring. After 24 hours of incubation, the subsequent bacterial growth was evaluated by taking the absorbance of the medium at 600 nm.^40^

#### Cytotoxicity Assay

By taking a 24-well plate, A549 cells were cultured for three days with approximately 2.5 × 10^4^ cells per well. The cells were then washed with PBS buffer having pH = 7.4, and then 500 μL of fresh DMEM media was added to each well. After that, different concentrations of CQDs and the drug-loaded CQDs (C-CQDs) were added and incubated for 24 hours in different plates. Followed by washing with PBS and addition of fresh DMEM media, freshly prepared 30 μL of stock MTT reagent (5 mg/mL) was added and kept for 4 hours. After 4 hours, the supernatant was carefully discarded, and the violet crystal of formazan was dissolved in 500 μL of DMSO. The absorbance was measured subsequently at 570 nm.^41^ The control was taken with 100% viability with the cells which are not treated with any sample.

#### Measurement of Quantum Yield

By taking Rhodamine-6G (R-6G) as the standard reference, and by incorporating the area under the fluorescence curve and absorbance values, the fluorescence quantum yield of the CQDs was calculated. From the UV-Vis spectra, the observed absorbance is less than 0.1 for both the standard and the sample. By diluting them with MILLI-Q water, both the sample and reference are taken. By using equation 14, the fluorescence quantum yield was calculated taking Rhodamine-6G as a standard, having a quantum yield of 95%.^42^

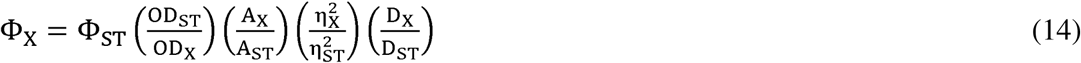

Where, Φ_x_ represents the quantum yield of CQDs, and Φ_ST_ represents the corresponding quantum yield of standard R-6G. OD_ST_ and OD_x_ are the standard and reference optical densities, respectively. The absorbances of the sample and standard are A_x_ and A_ST_ respectively. D_X_ and D_ST_ are the refractive index of the sample and reference, respectively. D_x_ and D_ST_ are the dilution ratios of sample and standard, respectively.^43^

## Results and Discussion

For the synthesis of CQDs, an easier, greener approach has been taken, with natural sources as a precursor and using reflux methods at 210 °C for 3.5 hours, followed by filtration and centrifugation, which results in a brown color homogeneous solution (details in the experimental section). Under 365 nm of ultraviolet radiation, a notable change in the color was observed, which turned into green (scheme 1). To get a powder form of CQDs, the formed brown residue was then dried at 60 °C by keeping it on top of the hot plate, followed by placing it inside a vacuum desiccator, and using a vacuum pump. To determine various solid-state characterization purposes, the powder form was stored at room temperature in a bottle wrapped in aluminium foil. The as-synthesized CQDs in their solution form in DI water were taken to characterize the optical properties of the sample, such as UV-Vis spectroscopy and fluorescence emission have two distinct peaks, one at 331 nm and the other at 284 nm. The *n* – *n*.* transition due to spectroscopy. Figure 1A (inset) represents the UV-Vis spectrum of the CQDs in water, which C=O bonds mainly results in a peak at 331 nm, and a wavelength region of 284 nm was found due to the C=C sp^2^ hybridized π –π* transition.^44–45^ However, we have found a broader peak ranging up to 530 nm, which leads to a range of electronic transitions. In Figure S2, the absorption spectra have been taken for only the extract in comparison with the CQDs. And we have found there is no such distinct peak in the case of only the extract. When we move from an excitation of 320 nm to 450 nm of wavelength, it has been found that the corresponding emission intensity decreases with a red shift (Figure 1A). Due to random distribution, increase in size, and presence of various functional groups over the surface, this trend in fluorescence spectra was observed.^46^ When a precise wavelength of light is anticipated, particles of equal size will be emitted, and particles of different sizes will be emitted with the change of excitation wavelength. Importantly, different band gaps will result due to the nonuniform particle size of CQDs. Along with that, the various functional groups also result in photon emission to come to the ground state via various routes because of the ability to generate their own energy levels. Using a 532 nm laser, the Raman Spectra of CQDs were collected to get the idea of the different types of structural characteristics of carbon, and we have found two major peaks for carbon; one at 1261 cm^-1^, which corresponds to the D-band of sp^3^ carbon atoms, represents the out-of-plane vibrations of sp^3^ hybridized carbon atoms. The peak at 1555 cm^-1^, which corresponds to the G-band of the sp^2^ carbon, represents the in-plane vibration of the sp^2^ hybridized carbon atoms.^47^ Notably, the intensity of the G-band is higher than that of the D-band, and the I_D_/I_G_ ratio was found to be 0.77, confirming the intrinsic, well-ordered carbon materials with symmetry-allowed vibration of sp^2^ carbons.^48–49^ However, the D-band confirms the presence of functional groups or defects on the surface, and to get an idea of that, FTIR spectra were recorded, and from the spectra, different types of functional groups were confirmed (Figure 1C). The presence of -OH groups was confirmed from the FTIR peak attributed to 3340 -3835 cm^-1^ due to the -OH stretching, and the peak in the range of 2690-3340 cm^-1^ is due to the C-H stretching.^50–51^ The carbonyl group present on the surface of the CQDs represents a peak in the range of 1700 cm^-1^.^52^ The C=C has the peak in the region of 1580 cm^-1^.^53^ In the fingerprint region, various peaks were found, and the peak at 955 cm^-1^ is due to the C-O bond.^54^ Because of the presence of various functional groups, it mainly helps with the solubility and adsorptivity of CQDs.^55^ In addition to that, by taking only the extract of the mixture, in the FTIR spectrum, we have found two additional peaks at 2312 cm^-1^ and 2364 cm^-1^ (Figure 1C), mainly due to unsaturated carbon-carbon bonds that may be lost after the formation of CQDs.^56^ By taking the excitation spectra from 300 nm to 550 nm, keeping emission from 310 to 700 nm, the 3D contour profile was taken, and the Bird-eye view is shown in Figure 1D. From the spectra, it has been found that there is only one major peak, with the corresponding excitation and emission wavelengths. To know the crystallinity and structure of green-synthesized CQDs, X-ray diffraction analysis was performed, as shown in Figure 1E. It was revealed from the diffraction pattern that at 2θ values of 20^°^ and 42^°,^ there are two major peaks which correspond to the plane (002) and (100), respectively.^57–58^ The crystallographic purity in the sample was confirmed due to no such spurious diffraction. To know the particle size distribution of the CQDs, FESEM analysis was performed and is shown in Figure 1F. It was confirmed that the formation of CQDs have a spherical shape with an average particle size of 9.25 nm (Inset, Figure 1F).

**Scheme 1.**
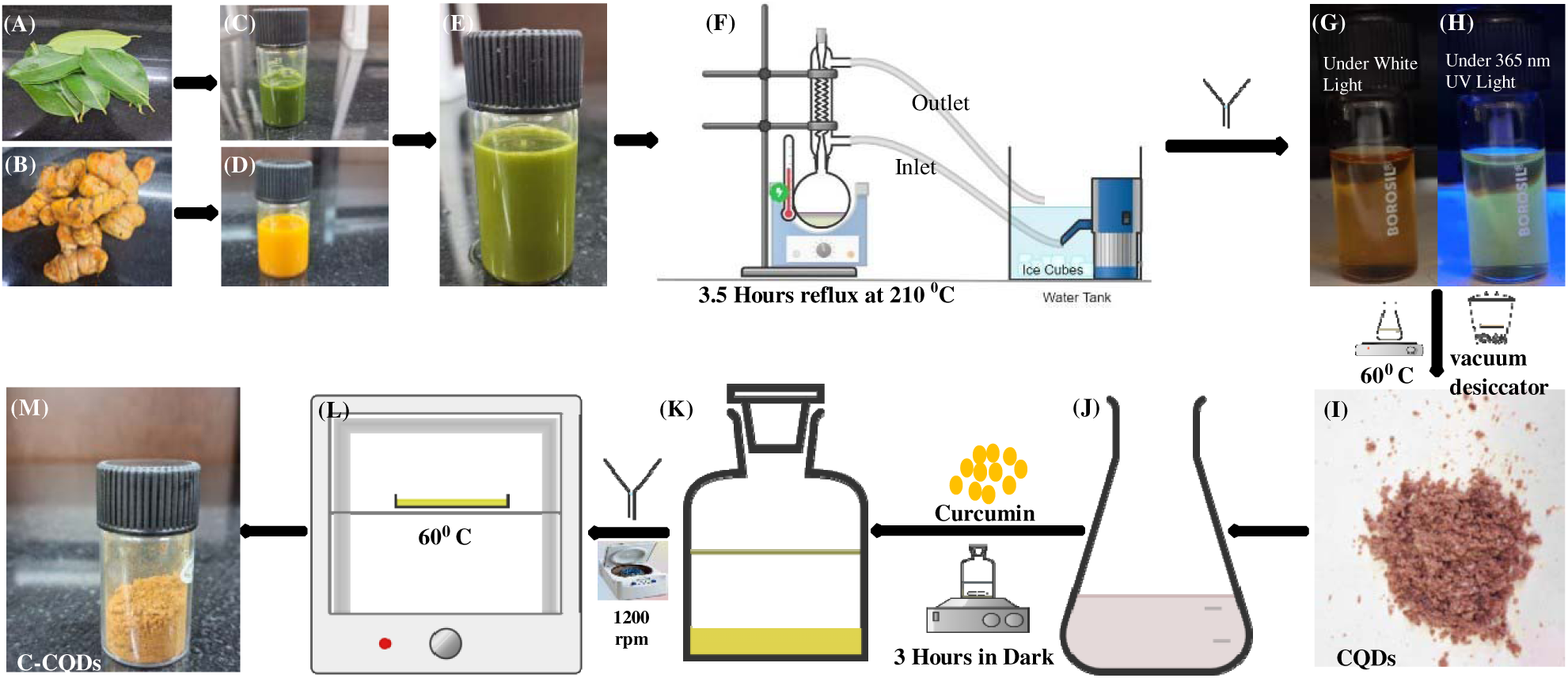
Green synthesis of Carbon Quantum Dots (CQD) and Curcumin loaded Carbon Quantum Dots (C-CQD). (A) and (B) The precursors taken for the synthesis of Carbon Quantum Dots (CQDs) are Jamun leaves and raw turmeric, respectively, from where (C) and (D) the juice extracts were collected. (E) A 3:1 mixture of corresponding juice extracts is taken for the green synthesis. (F) The setup used for the green synthesis of Carbon Quantum Dots (CQDs) using reflux conditions, which forms a brown color solution under normal light (G), turns green (H) under 365 nm UV light. Under 60 °C, on top of the hot plate, followed by a vacuum desiccator, which will form a solid form of CQDs (I). In its solution form with water (J), the addition of curcumin followed by a continuous magnetic stirring in dark conditions will result curcumin-loaded CQDs (K), which after filtration followed by centrifugation, results in C-CQDs that are kept in a hot oven (L) at 60 °C, resulting in solid C-CQDs (M)

**Figure 1.**
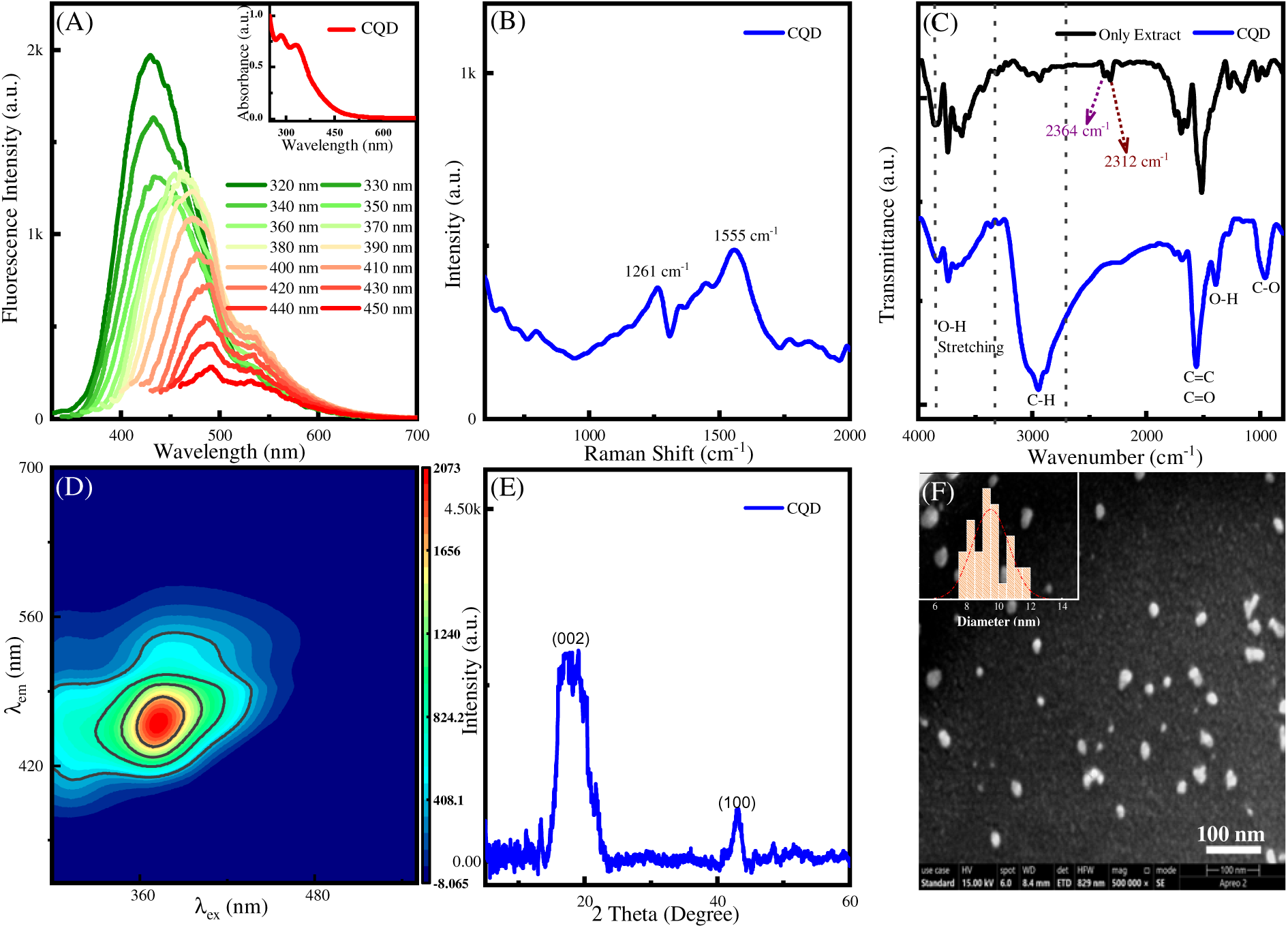
Characterization of CQDs. (A) Excitation wavelength-dependent emission spectra of CQDs at λ_ex_ *=* 320–450 nm. The inset shows the two peaks of UV-Vis peaks at λ_max_= 284 nm and 339 nm, (B) At 532 nm excitation wavelength, the Raman spectra of CQDs have been taken, (C) Fourier Transform Infrared (FTIR) spectra of CQDs compared with only extracts, (D) 3D Excitation dependent emission Spectrum (EEMS) of CQDs, (E) XRD spectra of CQDs shows one broad peak at 2θ = 20^0^ for (002) and a sharp peak at 42^0^ for (100). (F) FESEM image of CQD showing the average particle size of around 9.25 nm. Scale Bar = 100 nm

Emboldened by these results, we further tried to understand the detailed elemental compositional information and to examine the functional groups present on the surface of the CQDs and C-CQDs. X-ray photoelectron spectroscopy (XPS) was used to examine the surface-modified features of the CQDs and C-CQDs.^59^ In Figure 2, the XPS spectra were presented, which indicate the corresponding findings of CQDs and C-CQDs, including the functional groups and electronic state as well. In Figure 2A, the full XPS spectrum of CQDs is given, and it reveals the presence of Carbon (C) and Oxygen (O) atoms, mainly with the relative percentage of 75.34% and 23.16%, respectively, with some other trace elements observed in very negligible amounts. From the full spectrum, it has been found that Carbon (C1s) forms a sharp peak with a typical binding energy of 285 eV and Oxygen (O1s) with a binding energy of 533.5 eV, which are the characteristics of those elements.^60^ However, the core levels were seen with a low set of pass energies, and a broad peak was found for both Carbon and Oxygen at that particular range. The deconvolution of the C1s binding peak represents three major peaks (Figure 2B). The binding energy peak of the C1s is the strongest at 284.54 eV which is due to graphitic sp^2^ hybridised carbon confirmed from the Raman spectra as well (G-band).^61^ However, the peaks at 286.03 eV and 287.56 eV are due to C-O and C=O groups, respectively.^62^ The O1s spectrum (Figure 2C) upon deconvolution shows two peaks, one at 531.73 eV and another at 532.66 eV, corresponding to the C=O groups and C-O groups, respectively.^63^ Afterwards, we have also taken the XPS spectra of C-CQDs, and from the spectra, it has been found that both the Carbon and Oxygen peaks are present at the binding energies of 285 eV and 533.5 eV, respectively (Figure 2D). However, the relative percentage of Carbon and Oxygen is 77.69 and 19.71%, respectively. In comparison with the CQDs, the Oxygen percentage of C-CQDs decreased, and this may be because of some functional groups that might take part in the attachment with curcumin, or some small molecules may be lost after the attachment of curcumin.^64–65^ Upon deconvolution of the C1s spectra, it has been found that there are three different peaks at 284.61 eV, 286.02 eV, and 288.01 eV (Figure 2E). In case of C-C binding energy, which is at 284.61 eV, the intensity is not as much affected as compared to the intensity that has been found from CQD only. It may be because the sp^2^ hybridised carbon atoms, which are at the core, are not much affected by the attachment.^66^ However, the binding energy due to the functional groups (286.02 eV and 288.01 eV) shows a decrease in the intensity with a slight shift in the spectra towards the right. From the deconvolution of Oxygen (O1s) spectra, two peaks were found similar to those of CQDs. The peak at 531.67 is due to C-O groups, and the peak at 532.69 is due to C=O groups. As compared to CQDs, there is a slight decrease in the intensity of both peaks.

**Figure 2.**
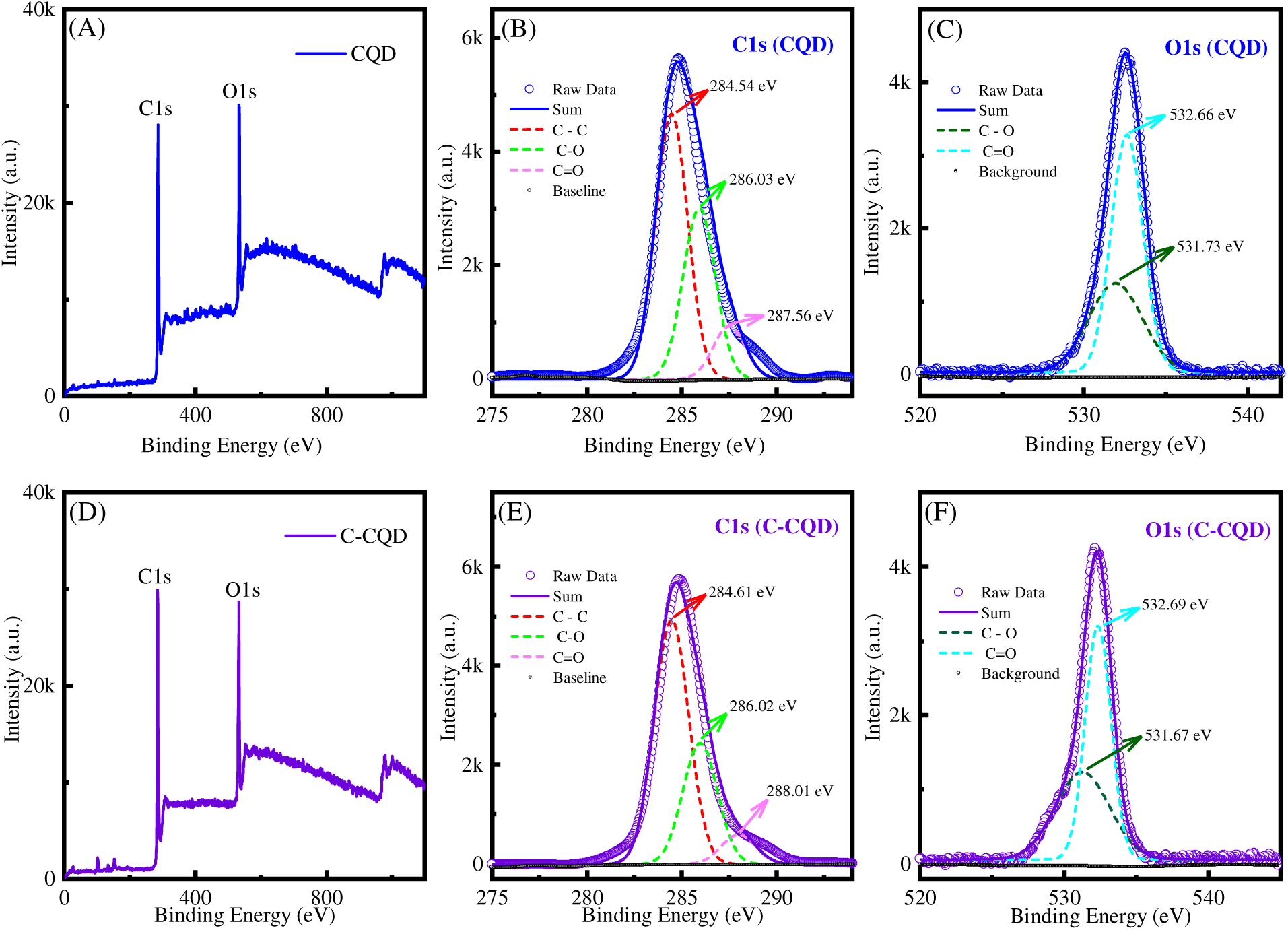
XPS spectra of CQDs and C-CQDs. (A) Total binding energy survey of CQDs and (B) the corresponding binding energy graph with C1s and (C) with O1s (C). (D) Total Survey of C-CQDs and (E) the corresponding binding energy graph with C1s and (F) O1s.

Additionally, some important parameters like hydrodynamic size distribution, zeta potential, and electrophoretic mobility can be studied by using dynamic light scattering (DLS) techniques (Figure 3).^67^ Using a Zeta-sizer, the average hydrodynamic diameter of the CQDs was found to be 18.66 nm, with the distribution from 11.23 nm to 38.56 nm(Figure 3A). However, it is found that the average particle size of the CQDs was larger than that found from FESEM (i.e., 9.25 nm), as the hydrodynamic diameter of the particles in water is larger than only CQDs measured in a vacuum.^68^ As the particles in their vacuum state are of different sizes, so the water that is surrounded by that particle is also in different ratios, which results in a larger hydrodynamic diameter than bare CQDs. Comparatively, to know the hydrodynamic diameter of C-CQDs, we have measured the hydrodynamic diameter of C-CQDs, and that was found with an average diameter of 59.07 nm, ranging from 43.125 nm to 63.03 nm (Figure 3G) confirms that the average diameter of C-CQD is larger than that of only CQDs because with the attachment of Curcumin on the surface, various distribution of the size was found. Notably, without any functional group on the surface, CQDs can easily precipitate in aqueous-based solvents, and on the other hand, multiple functional groups on the surface improve the dispersion in water-based solvents, which in the long term, is vital for the colloidal stability and good dispersal in a water-based solvent.^61, 69^ Additionally, to know the surface charges that exist on the surface, the zeta potential measurement was done, and the average zeta potential was found to be -13.92 mV with 24.65 mV of peak width (Figure 3B). The results indicate that the presence of anionic groups or ions, such as -COOH, that are on the surface of CQDs provides the overall negative charge on the surface.^70^ In response to the applied electric field, the -vely charged CQDs moves, and one of the crucial parameters to measure that is electrophoretic mobility, and the mean value was found to be -1.26 Cm^2^/Vs (Figure 3C) and due to this small value, the CQDs move very slowly under the influence of the electric field. The -ve value of both zeta potential and mobility indicates the surface negative charge, and as the particles in their vacuum state are of different sizes, so the water that is surrounded by those particles is also in different ratios, which results in a larger migration towards the positively charged electrode.^71^ The elemental composition and spatial distribution of the CQDs can be explored by using a scanning electron microscope equipped with energy-dispersive X-ray (EDX) analysis. From the EDX micrograph, it has been found that the majority of the elements present are Carbon (C) and Oxygen (O), with atomic percentages of 79.17 and 20.83, respectively (Figure 3D), and their spatial elemental distribution was given in Figures 3E and 3F. The functional groups that present on the surface helps for the adsorption of foreign materials. By using equation 9, the curcumin adsorption efficiency was calculated and was found to be 34%.

**Figure 3.**
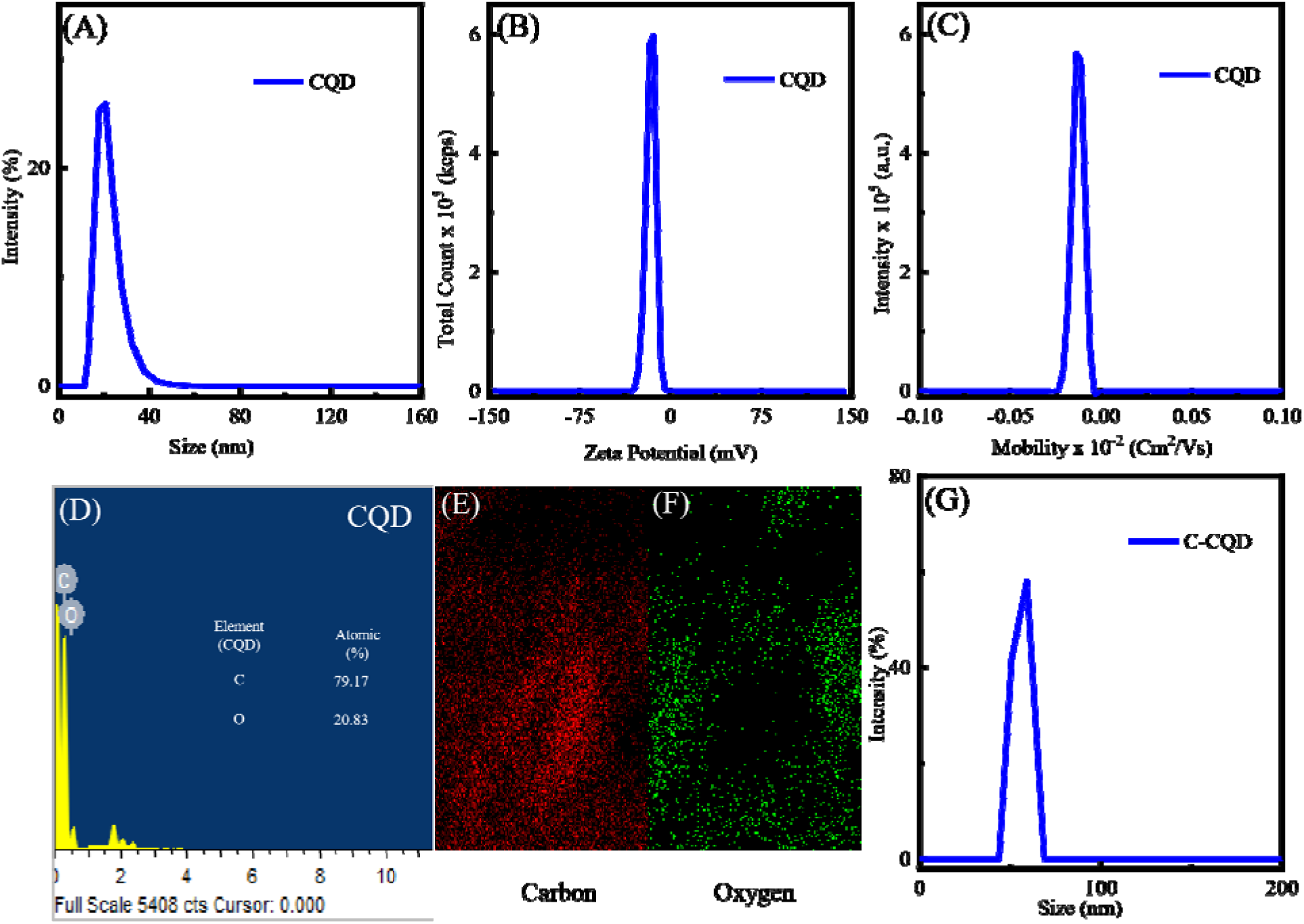
DLS Spectra and EDX micrograph of CQD and C-CQD. (A) Dynamic light Scattering experiment with CQD confirms the hydrodynamic diameter with a peak at 18.66 nm. (B) Zeta potential of CQDs showing a peak at -17.23 mV, ranging from -33.04 mV to -4.58 mV, with (D) electrophoretic mobility of -0.0001102 Cm^2^/Vs. (A) EDX spectra of CQDs confirm the major elements present are Carbon (C) and Oxygen (O). Elemental mapping of CQDs, (E) for carbon and (F) for Oxygen. The inset table displaying the weight percentage and atomic percentage of CQDs. (G) The hydrodynamic diameter of C-CQD, with a peak at 59.07 nm, confirms a higher hydrodynamic diameter than CQDs.

Next, we have explored the synthesis of C-CQDs, and in the experimental section, the detailed synthetic procedure of C-CQD was discussed. After the synthesis of C-CQDs, it was further characterized by using various Spectroscopic techniques. In Figure 4A, inset, the UV-Vis spectra of C-CQD were shown in comparison with those of CQDs. Including those two small hump that are there in CQDs as well (284 nm and 334 nm), another hump at 433 nm along with broad signal was found supporting the attachment of curcumin on the surface of CQDs.^72^ The as-synthesized C-CQD was found to be soluble in water, which is not generalized to the curcumin, which itself is not soluble in water. By taking the excitation wavelength from 380 nm to 500 nm, the corresponding fluorescence intensity was taken, and it was found that, like CQDs, C-CQDs were also showing excitation-dependent emission spectra with a decrease in intensity and red shift as well (Figure 4A). This trend in fluorescence spectra is due to the random distribution of size as well. Including the peak at 488 nm, another peak has been found at 535 nm, which may be due to the curcumin that is attached to the CQDs. So, the indication of two high-intensity peaks at 488 nm and 535 nm confirms the attachment of curcumin to CQDs. The 3D excitation-dependent emission matrix spectrum has been taken by taking excitation from 300 nm to 600 nm, keeping the emission range from 310 -700 nm. The bird-eye view of the spectra is given in Figure 4 B. Just like in Figure 4A, we can observe two peaks, respectively for CQDs and curcumin that are conjugated to CQDs. Inside the contour, a broad black straight line is observed, and that is due to the Rayleigh scattering, where λ_ex_ = λ_em_.^36^ Despite these results, in terms of functional groups, a difference in the signals was found due to C-CQDs when comparing with CQDs and Curcumin only. In Figure 4C, a broad peak from 3255 cm^-1^ to 2879 cm^-1^ was found due to the formation of intermolecular H-bonding of curcumin with CQDs.^73^ Similarly, another broad peak from 1567 cm^-1^ to 981 cm^-1^ was found due to C=O stretching vibration that is coupled with the other groups.^73^ For only curcumin, due to the presence of phenolic -OH, its stretching vibration results in a peak at 3452 cm^-1^. Additionally, the peak at 1631 cm^-1^ corresponds to the C=C peak, and for the ketonegroup and C-O groups, peaks at 1465 cm^-1^ and 1162 cm^-1^ were found.^74–75^ Figure 4D represents the Raman spectra of C-CQDs in comparison with CQDs and curcumin. In C-CQDs, the peaks, which are at 1263 cm^-1^ and 1561 cm^-1^, are due to the D-band and G-bands of carbon. For the benzene ring, the peaks at 1598 cm^-1^ and 1647 cm^-1^ are found in the case of curcumin. In case of C-CQDs, for benzene, both the peaks are found at 1563 cm^-1^ and 1645 cm^-1^.^76^ The peak at 1101 cm^-1^ in the case of curcumin is due to the aromatic bending, which is also found in the case of C-CQDs at 1107 cm^-1^. Additionally, the peak at 1780 cm^-1^ in C-CQDs is due to the C=O stretching band, which is only found in the keto form of curcumin and not in the enol form.^72^ So, the peak exists due to the attachment of curcumin with CQDs.

**Figure 4.**
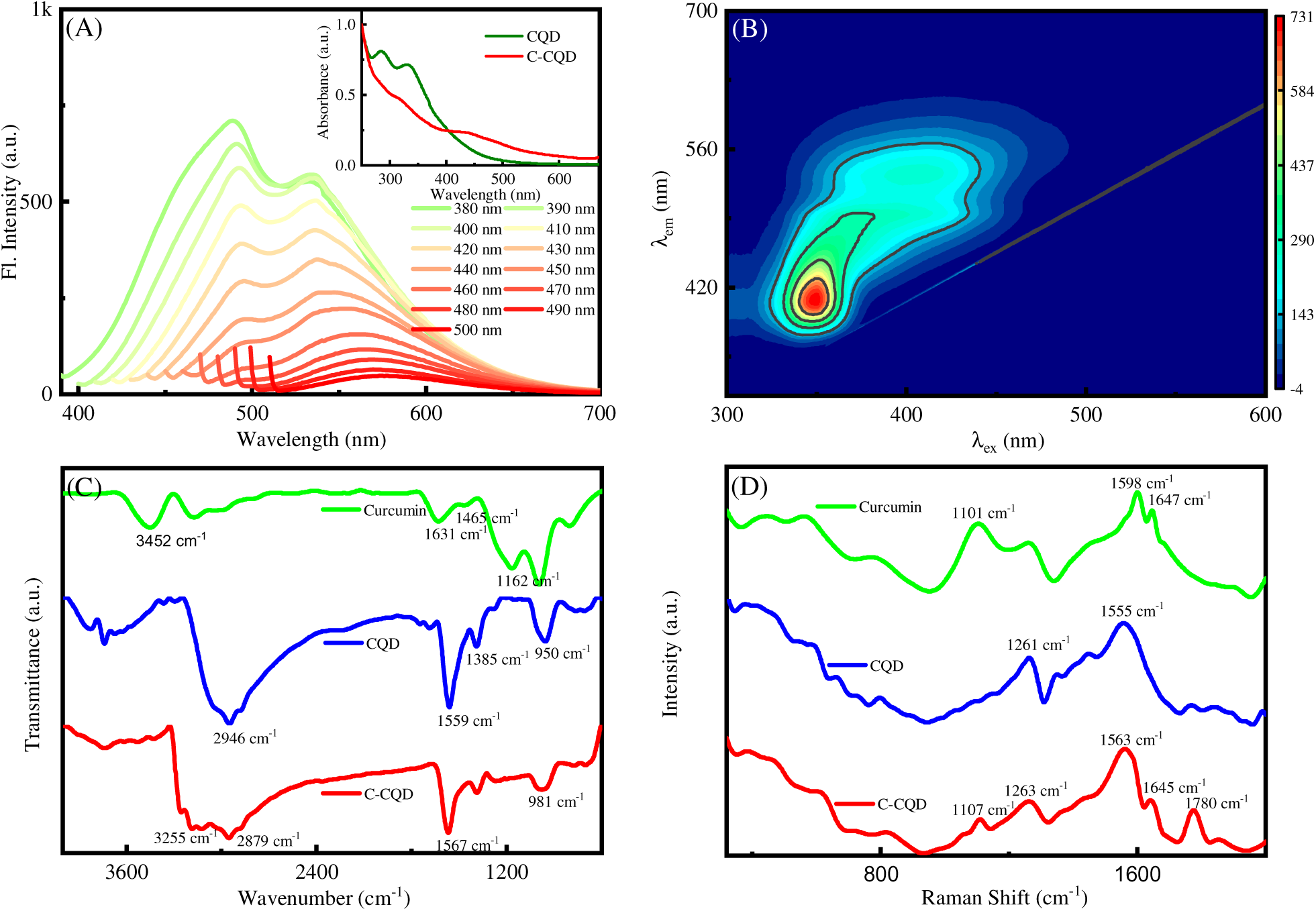
Characterization of Curcumin loaded CQDs (C-CQDs). (A) Excitation wavelength-dependent fluorescence emission of C-CQDs with excitation wavelengths from 380 nm–500 nm. The inset displays the absorption spectra of C-CQDs and it’s comparison with that of only CQDs (B) 3D Excitation-Emission Matrix Spectrum of C-CQDs. (C) Fourier Transform Infrared (FTIR) spectra of C-CQD in comparison with CQD and curcumin. (D) Raman spectra of C-CQD in comparison with CQD and curcumin, excited at 532 nm excitation wavelength.

With curcumin conjugated on the surface of CQDs, the binding interaction of C-CQDs with serum albumin protein, BSA, can be of interest. The binding and interaction depend on several factors, such as noncovalent, hydrophobic, and nonspecific, as well.^77^ However, the nanocrystal size, curvature, and surface chemistry are the factors on which it depends the most,^78–79^ and we are interested in knowing the spectroscopic understanding of the binding of protein such as Bovine Serum Albumin (BSA) with the hydrophobic drug curcumin after the attachment with CQDs; that is C-CQDs, which is our system of interest. So, we have conducted a series of spectroscopic measurements, such as steady-state and time-resolved fluorescence measurements. With the interaction of the BSA protein with a fluorescence quencher (C-CQDs), the quenching of fluorescence of a fluorophore can be attributed to various processes, like ground-state complex formation or excited-state energy transfer.^77^ Based on that mechanism, there are two different types of fluorescence quenching: static and dynamic. Here, we have studied the strength of interaction between C-CQDs and BSA. At 298 K, the steady state fluorescence spectra of BSA were taken with sequential addition of C-CQDs from 0.5 μL to 22 μL from the stock solution (0.25 μg/mL to 11 μg/mL) at 280 nm excitation. Comparison with the fluorescence spectra of only BSA, which is maximum at λ_em_ = 338 nm, the fluorescence intensity of the mixture decreased with the addition of C-CQDs (Figure 5A) at that emission maximum. However, in the case of only CQDs, no quenching occurs with the addition of CQDs (Figure S3). To explore the molecular route involved in the quenching of fluorescence during the interaction of C-CQDs with BSA, the familiar Stern-Volmer equation (SV) was used. Using equation 4, the Stern-Volmer quenching constant was calculated and was found to be 0.14 (Figure 5B). The Stern-Volmer quenching constant, K_SV_, typically tells the quenching process’s efficiency. Using equation 5, the number of binding sites per protein was obtained as 1.06, given in Figure 5C. In Figure 5C, the fluorescence emission decays of the BSA-[C-CQDs] are shown. It has been found that the fluorescence lifetime of BSA decreases gradually with the addition of more C-CQDs (Figure 5F). Furthermore, the 3D excitation-emission matrix spectrum was studied by taking emission from 210 nm to 650 nm, keeping the excitation from 200 nm to 400 nm for both BSA and BSA with C-CQDs. In Figure 5D, the bird-eye view of 25 μM of BSA was given, where only one peak was found with emission maximum at 338 nm at excitation of 280 nm. However, after the addition of C-CQDS, in addition to the BSA peak, two other peaks at 465 nm and 460 nm were found, respectively. Both peaks correspond to the characteristic peak of C-CQDs. The bird-eye view of the 3D excitation-emission matrix spectrum is shown in Figure 5E. From the decay profile, two different decay components were found. One with a shorter lifetime component of 2.35-1.83 ns, having an amplitude of 9-16%, and the longer component with a lifetime of 5.97-3.84 ns, with an amplitude of 91-84%, after the gradual addition of C-CQDs at 298 K. The longer lifetime (6.0 ns) may be associated with free BSA, whereas the obtained shorter components may be due to the dynamic quenching of BSA-C-CQDs.^77^ For the calculation of the average lifetime of the corresponding components, equation 7 has been used, and it has been found that the average lifetime of the components decreases from 5.644 to 3.52 ns with the increase of volume of C-CQDs from 5 to 20 μL (2.5 μg/mL to 10 μg/mL). The dynamic nature of the quenching process is further supported by the dynamic quenching process. The donor-acceptor overlap spectra in their normalised form of 25 μM BSA (Donor) and 2 μL (1 μg/mL) C-CQD (acceptor) are shown in Figure S1. When excited at 280 nm, the fluorescence intensity of the donor (25 μM BSA) got quenched, and the fluorescence intensity of the acceptor (C-CQDs) rises in the mixture as shown in Figure 6. Figure 6 clearly indicates the transfer of energy from the donor to the acceptor. We can observe that the donor fluorescence intensity decreases after the addition of acceptor (C-CQDs), and the energy transfer efficiency (E) from the donor to the acceptor was determined with respect to the increase in acceptor intensity rather than the quenching of the donor intensity. The intensity of the energy transfer was calculated by using equation 8 and was found to be 33%. To investigate the uptake mechanism of CQDs and C-CQDs inside the cancerous cells, the precultured glass-bottom petri dishes, each with a small cut glass slide of approx. 18 × 18 size, were kept ready as discussed in the experimental section. At first, to observe the Quantum dots (CQDs and C-CQDs), we have cells that were treated with CQDs, C-CQDs, along with a control where no quantum dots were added. Figure 7 A1 and A2 show the cells where no treatment was done. It was found from the image that no pores on the surface when there is no such treatment of quantum dots. However, when cells are treated with the DAPI Channel GFP Channel Merged CQDs, quantum dots have been found with small pores on the surface of the cells (Figure 7B1 and B2), and comparatively bigger pores have been found in cells treated with C-CQDs (Figure 7C1 and C2) with particles on the surface. These results suggest that, upon the treatment of quantum dots, the surface of the cells is affected mainly with pore formation,^80^ and there is a possibility that both the CQDs and C-CQDs are going inside the cell because of their minute sizes.^81^ To investigate the excellent biocompatibility and fluorescence properties of CQDs and C-CQDs, we have checked the imaging-based intracellular uptake of the CQDs and C-CQDs. For that, we have cultured cells in two different Petri dishes with A549 cells on the surface of an 18 × 18 glass cover slip that was placed inside the Petri dish with cell culture media (details given in the experimental section). In one Petri dish, we have treated the cells with both DAPI and CQDs and were imaged by using an EMCCD camera at two different channels (i.e., DAPI channel and GFP channel) using a halogen lamp as the excitation source with a 100X objective. Similar methods were used to check the cell imaging with C-CQDs as well. Figures 8A1 and A2 represent the fluorescence image of cells in the DAPI channel for CQDs and C-CQDs, respectively, where only the nucleus shows blue fluorescence due to the DAPI, as DAPI is selective only for the nucleus. However, with that EM gain and exposure time, both CQDs and C-CQDs do not show any imaging of the nucleus, though both have absorbance in the same region. It may be because of the very weak selective penetration of both dyes into the nucleus. Figures 8 B1 and 8 B2 are the fluorescence images in the GFP channel of that same cell with the same position that was treated with CQDs and C-CQDs, respectively. From the fluorescence images, only the cytoplasm area shows fluorescence due to CQDs and C-CQDs, respectively, or we can say that on treatment with quantum dots (CQDs and C-CQDs), it can enter the cytoplasmic area selectively. Figures 8 C1 and C2 are the merged images of both the DAPI and GFP channels.

**Figure 5.**
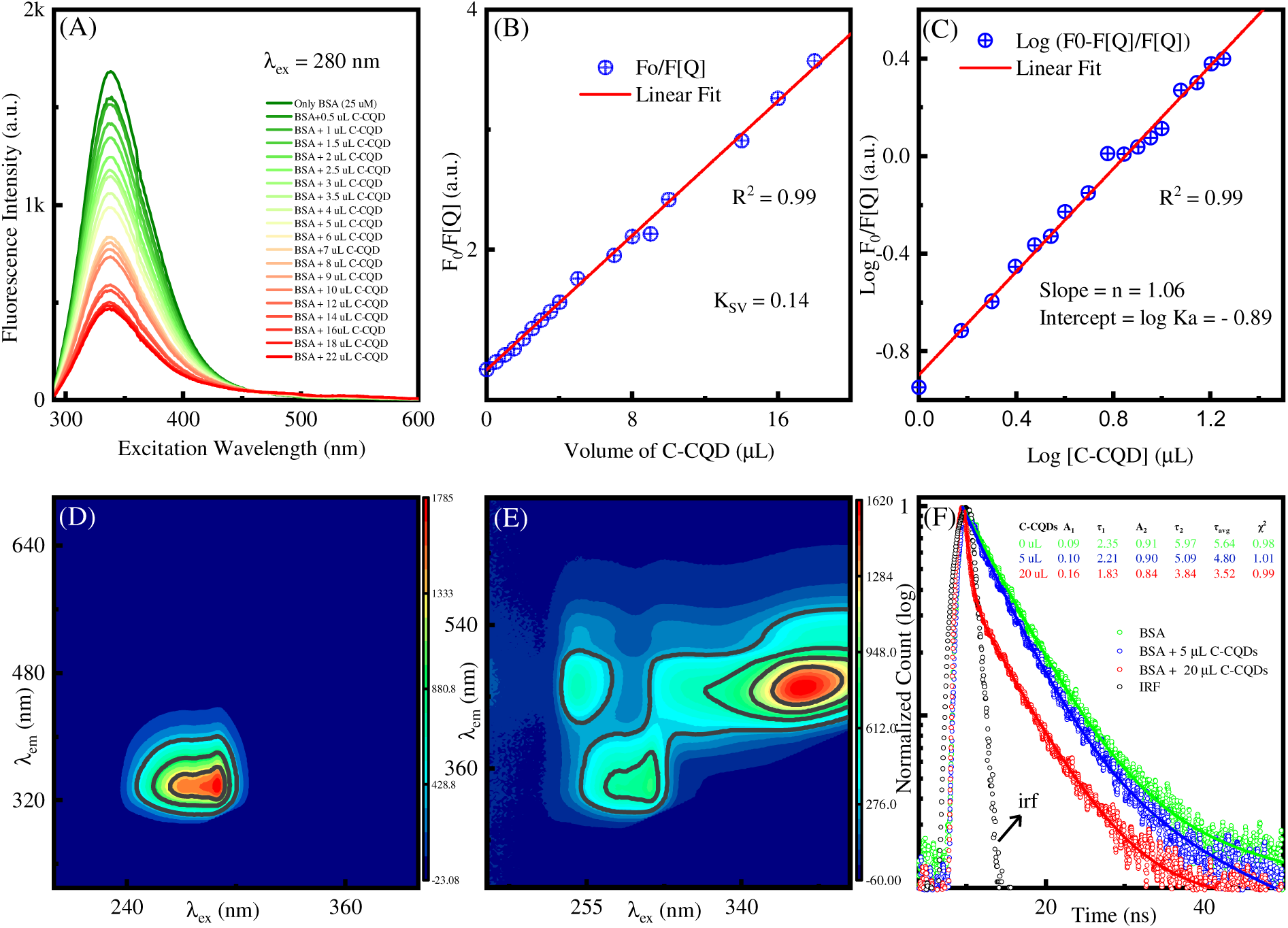
Effects of Bovine Serum Albumin with C-CQDs. (A) Steady-state fluorescence spectra of BSA upon the addition of C-CQDs at room temperature, (B) with corresponding Benesi-Hilderbrand plots showing the linear relationship to calculate the binding constant, and (C) the number of binding sites. (D) 3D Excitation-Emission Matrix Spectrum (EEMS) of BSA only, and (E) BSA with C-CQDs. (F) Fluorescence emission intensity decays of BSA along with C-CQDs addition.

**Figure 6.**
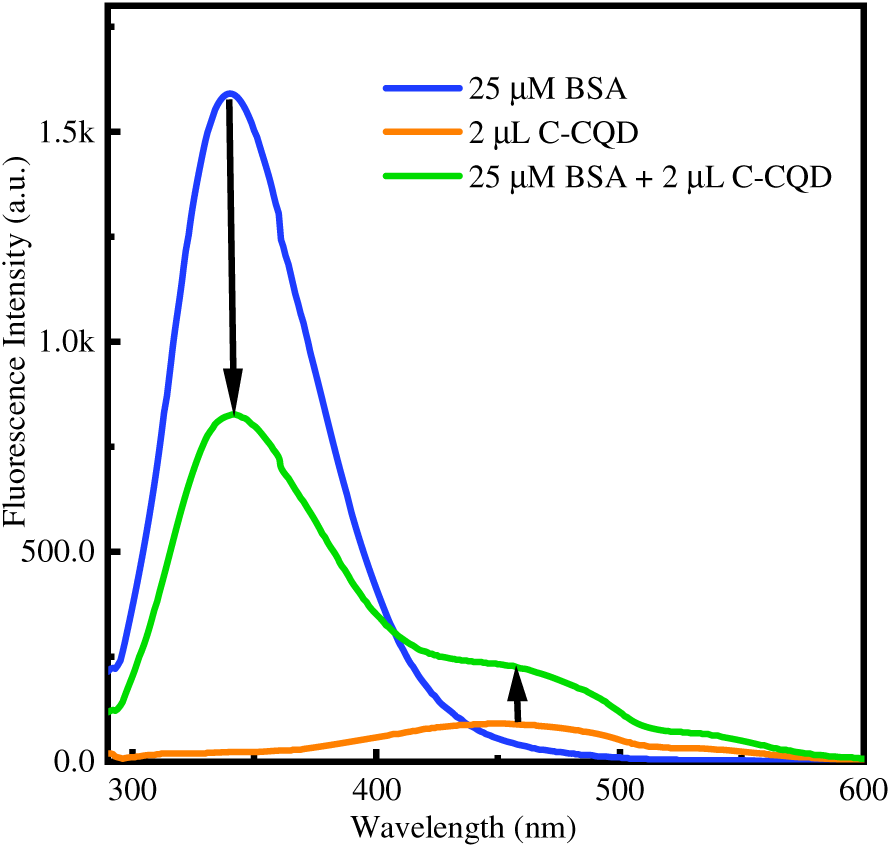
Fluorescence Resonance Energy Transfer (FRET) from BSA (25μM) to C-CQD at λ_ex_= 280 nm. From the energy transfer, it was confirmed that in the partially unfolded confirmation of protein BSA, the C-CQD binding happens with a drop in fluorescence intensity of BSA.

**Figure 7.**
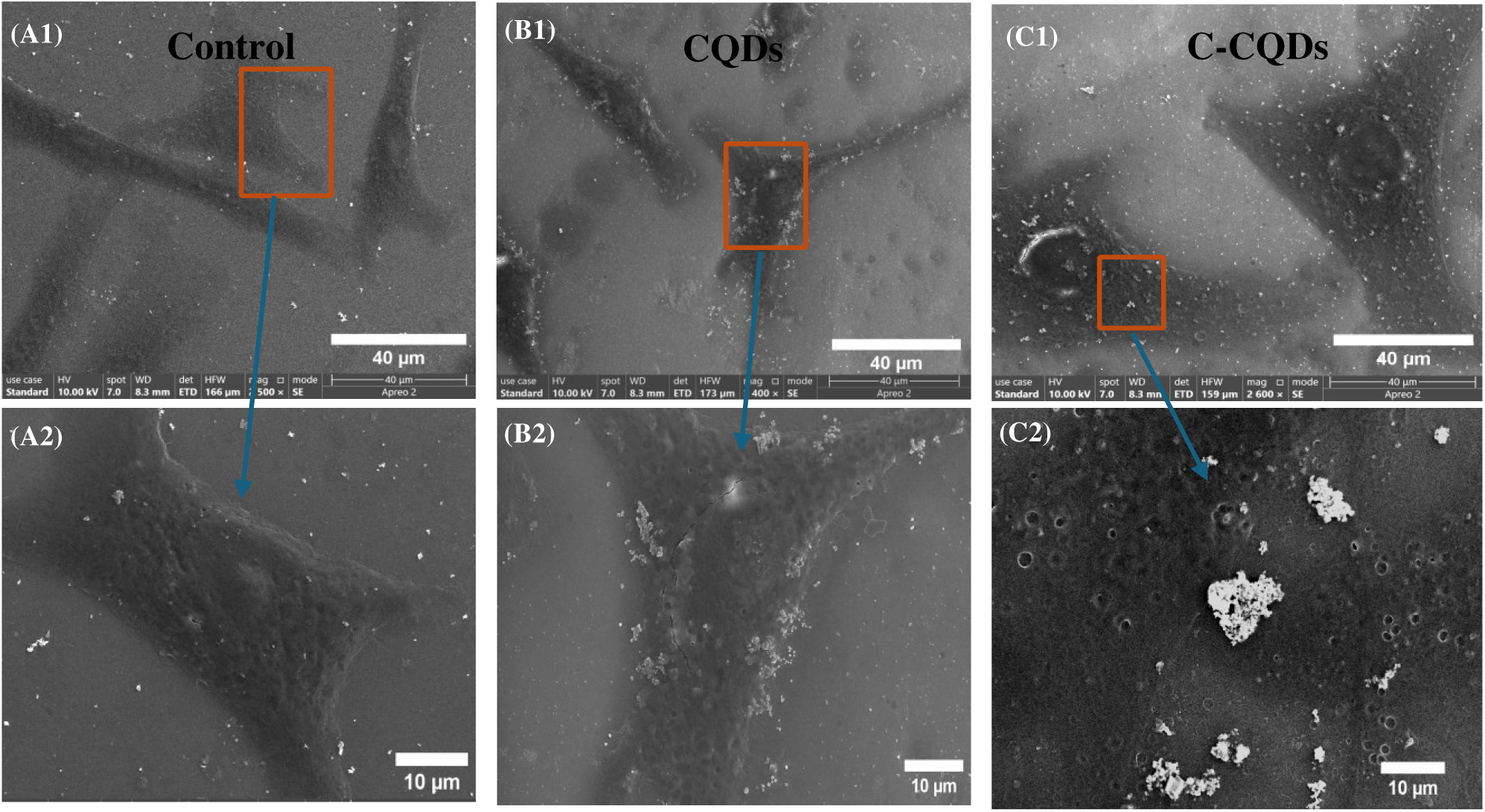
FESEM Image of A549 cells labeled with CQDs and C-CQDs. After incubation of cells with CQDs and C-CQDs in different cover glasses for 30 minutes, it was then fixed with 4% paraformaldehyde. (A1) and (A2) Cells without treatment of any sample were considered as a control, where no pore formation was found on the cell membrane. (B1) and (B2) Treatment with CQDs shows the attachment of CQDs on the surface of the cell membrane, along with small pores at the cell membrane. (C1) and (C2) Cells treated with C-CQDs also confirm the primary attachment of C-CQDs with larger-sized, frequent larger pores on the surface. Scale bar of A1, B1 and C1 = 40 μm & Scale bar of A2, B2 and C2 are 10 μm.

**Figure 8.**
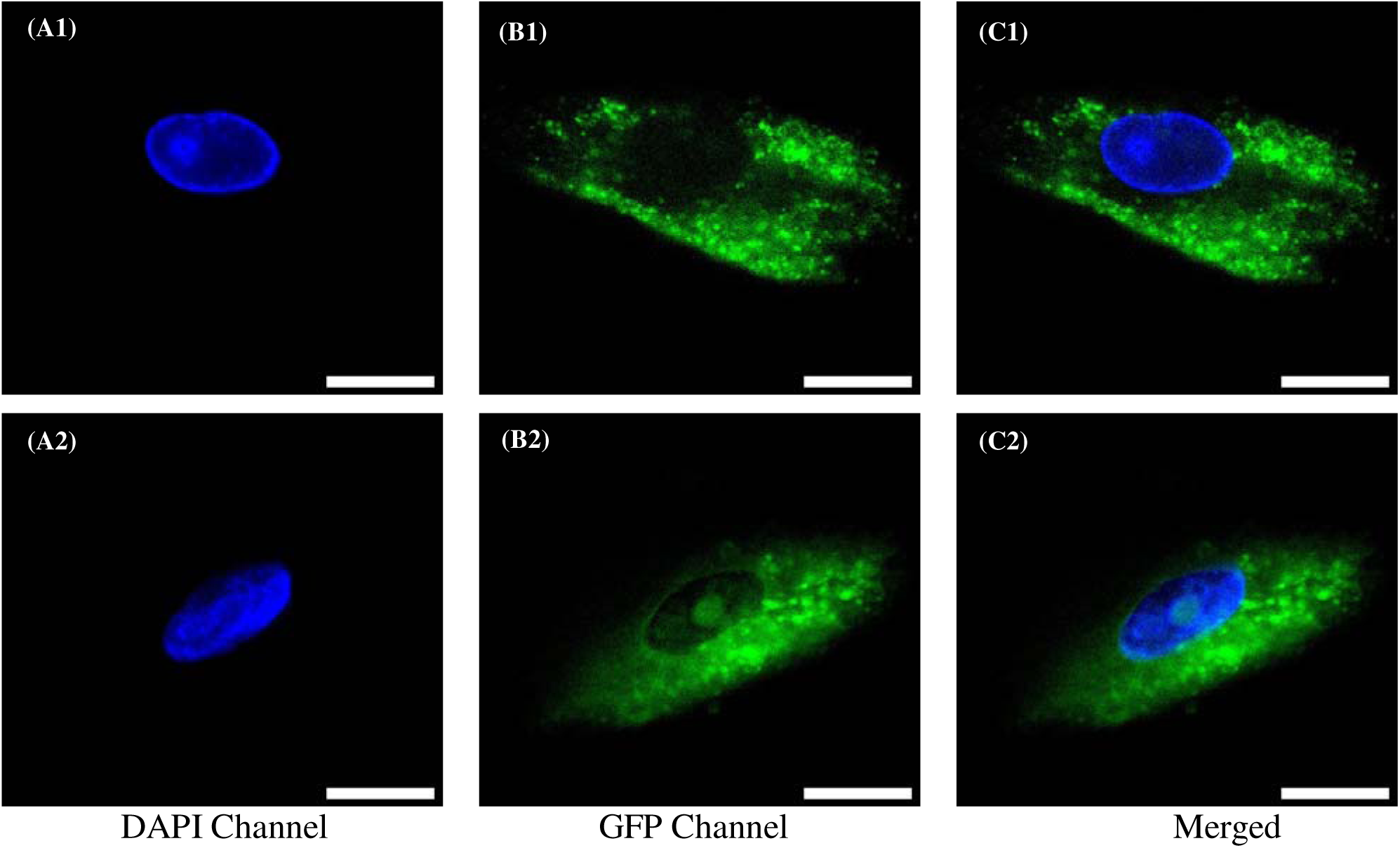
Fluorescence Imaging of the live A549 cell using carbon quantum dots (CQDs) and curcumin-loaded carbon quantum dots (C-CQDs) in different channels to confirm the penetration of CQDs and C-CQDs into the cytoplasm region selectively. (A1) Imaging of an A549 cell treated with DAPI and CQDs in the DAPI channel and (B1) the same cell in the GFP channel (C1) along with the merged of these two channels. (A2), (B2), and (C2), respectively, are the fluorescence images of A549 cells treated with C-CQDs and DAPI in the DAPI channel, GFP channel, and Merged (respectively). Scale Bar = 20 μm

Thereafter, we have taken the Confocal images of the live cell A549 after the incubation of CQDs and C-CQDs, separately considering the selective penetration of the quantum dots into the cytoplasm. In Figures 9A1 and A2, the confocal images were taken by treating CQDs and C-CQDs, respectively, with the laser excitation source of 405 nm, where the emission bandwidth was fixed at 430-470 nm. It has been found that both CQDs and C-CQDs show fluorescence, and we have obtained a fluorescence image in that excitation and emission range. Keeping the cell at the same position, we have taken the fluorescence image in the next channel, with the excitation source of 488 nm, and the emission was fixed at 500-550 nm (Figure 9B1 and B2). From the result, it has been found that the green fluorescence image was found only in the case of C-CQDs, wherea for CQDs, there is no fluorescence image of the cell. It may be because there is no efficient absorption and emission of CQDs in that range. However, in the case of C-CQDs, it shows fluorescence because of the significant absorption and emission in that particular range, and that is because of the curcumin that is attached to CQDs. Figures 9C1 and C2 are the brightfield images after treating CQDs and C-CQDs, respectively. Figures 9D1 and D2 are the merged images of all the channels. As we are getting the confocal images for the excitation-emission range of 488 nm / 500-550 nm only in the case of C-CQDs, we have taken the confocal image of the cells at different time periods of incubation to know the penetration of the drug efficiency at different times (Figure 10). It has been found that after 5 minutes of incubation, the drug started to enter the cell membrane, as shown in Figure 10 B. With the increase in the time of incubation from 5 minutes to 30 minutes, the fluorescence intensity increases (Figure 10 (A -E)), which may be due to more and more C-CQDs going inside the cell through the cell membrane due to the small particle size of C-CQDs.^81^ However, no fluorescence intensity was found in the control image (Figure 10A) as no C-CQDs were added to the cells. By using CQDs and C-CQDs as a fluorophore, we are eager to implement the microscopy imaging-based techniques to study the diffusion of those fluorophores in the plasma membrane of A549 cells in the cytoplasmic area. Using raster image correlation spectroscopy (RICS)techniques, which is a non-invasive and imaging technique to measure the diffusion coefficient of the fluorophore, we have measured the diffusion coefficient. The cells were seeded separately with CQDs and C-CQDs and imaged in a confocal laser scanning microscope using a 100X objective (details are explained in the experimental section). A minimum of four measurements were taken in the different regions of interest (ROIs) in the same cell, as shown in Figure 11 A and B. In Figure 11A, the cells were treated with CQDs, an image was taken, and within that cell, different ROIs were chosen. and then within those ROIs, the stack of images was taken at the same focal plane and area. In each ROI, a stack of 100 images has been taken for the analysis. Each of the stacks has a frame of 256 × 256 pixels. Figures 11 A1 and A2 are the stacks of images at different ROIs in the cytoplasm. After getting the corresponding 2D auto-correlation function (Figure 11 A3 and Figure A4), the diffusion coefficient was measured using the simFCS software discussed in the experimental section. The 3D representation with residues is given in Figures 11 A5 and 11A6. From the diffusion coefficient measurement, the diffusion coefficient was found to have an average value of 5.42 ± 0.16 μm^2^/sec. Similarly, the image has been taken in the case of cells that were treated with C-CQDs (Figure 11B). After choosing different ROIs in the same cell, a stack of 100 images was taken in each ROI. The stack of images is shown in Figure 11 B1 and B2. Each stack has 100 frames with a frame size of 256 × 256 pixels. After the analysis was done by using SimFCS software, the 2D autocorrelation function was represented in Figure 11 B3 and B4, and the corresponding 3D representations are given in Figure 11 (B5) and B6. From the diffusion measurement of C-CQDs, the diffusion coefficient was found to have an average value of 2.57 ± 0.14 μm^2^/sec. As per the Stokes-Einstein law, the diffusion coefficient (D_t_) value decreases with the increase in the hydrodynamic diameter. From the DLS measurement, the hydrodynamic diameter of C-CQDS is found to be larger than that of CQDs, so C-CQDS is found to have a slower diffusion than CQDs. The autocorrelation function of both CQDs and C-CQDs is given in Figure 12 A. From the figure, by taking the correlation factor (G0) along the Y axis with respect to pixel shift along the X-axis, it has been found that the decay of CQDs is faster than that of C-CQDs. The D_t_ values of the corresponding are shown in Figure 12(B). The diffusion coefficient of CQDs is found to be around 2-fold larger than that of C-CQDs.

**Figure 9.**
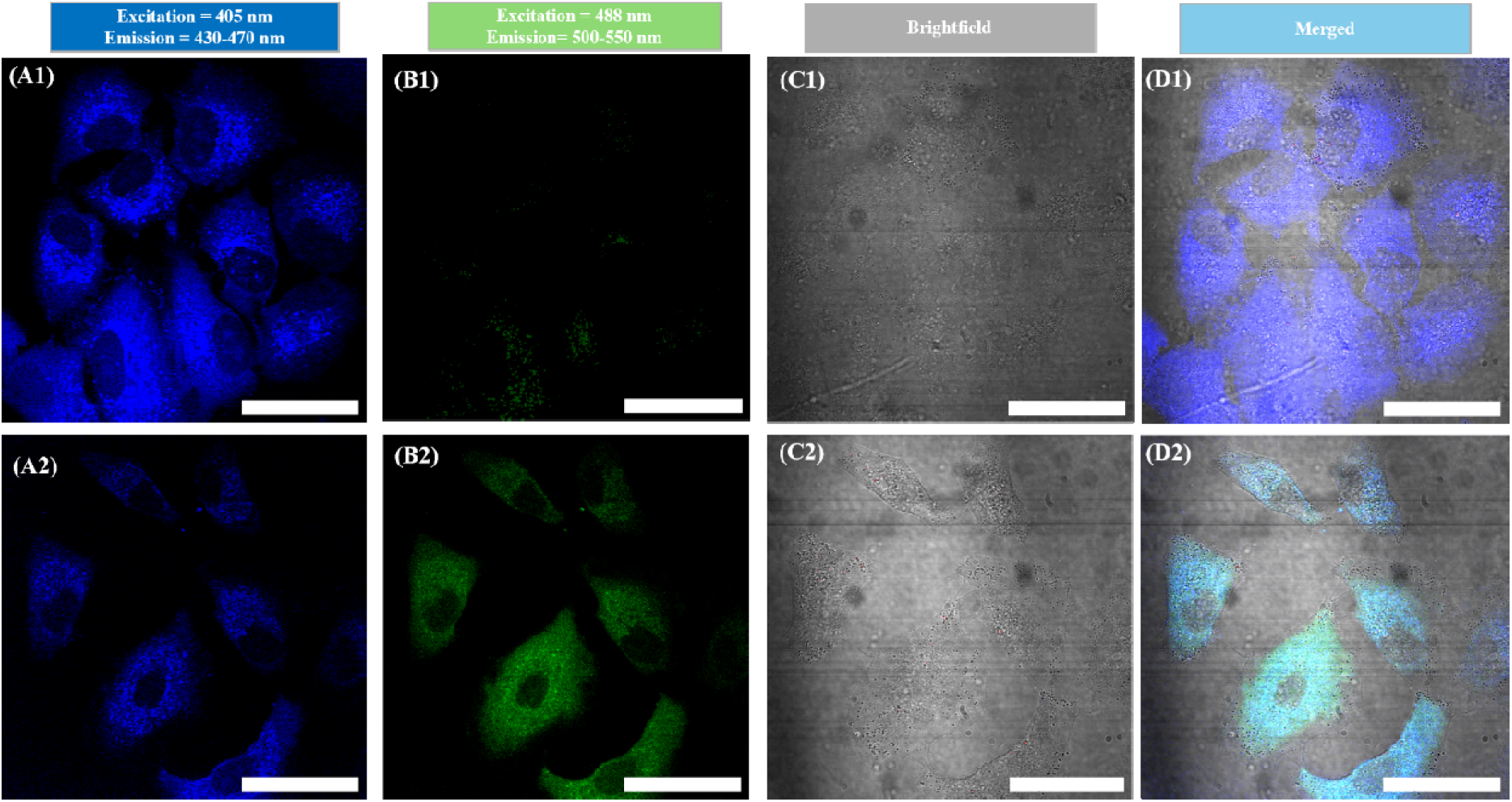
Confocal Image of the live A549 cell using carbon quantum dots (CQDs) and curcumin-loaded carbon quantum dots (C-CQDs). (A1, B1, C1, and D1) are the fluorescence images of live A549 using CQDs, and (A2, B2, C2, and D2) are for curcumin-loaded carbon quantum dots (C-CQDs). A1 and A2 have excitation and emission at 405 nm, 450 ± 20 nm, respectively. B1 and B2 have excitation and emission at 488 nm, 525 ± 25 nm, respectively. C1 and C2 are the transmitted light emissions. D1 and D2 are the merged images of all the channels. Scale Bar = 40 μm

**Figure 10.**
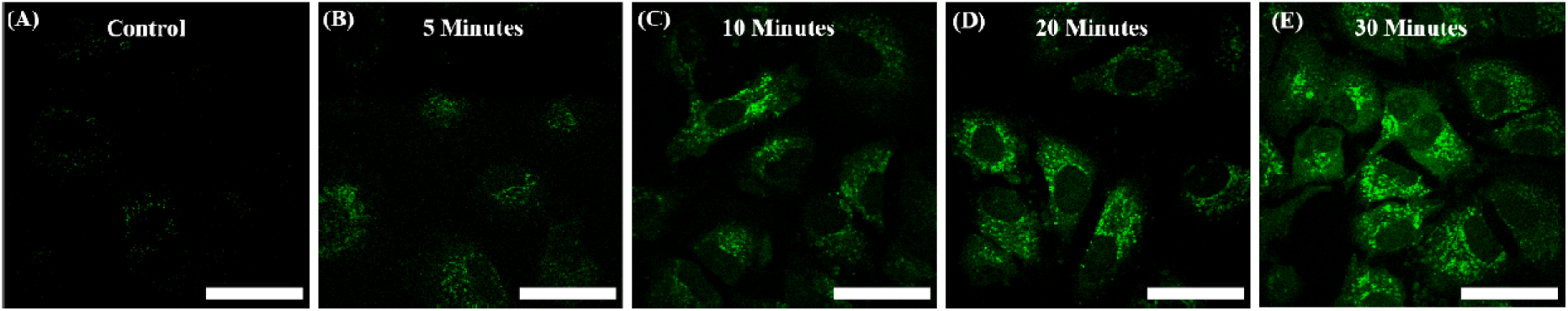
Time-dependent cellular uptake study. The excitation and emissions are 488 nm and 525 ± 25 nm, respectively. (A) Control images of A549 cells without the addition of C-CQDs. (B) Confocal image of A549 cells after 5 minutes, (C) after 10 minutes, (D) after 20 minutes, and (E) after 30 minutes. The results show an increase in the fluorescence intensity with an increase in incubation time. Scale Bar = 40 μm.

**Figure 11.**
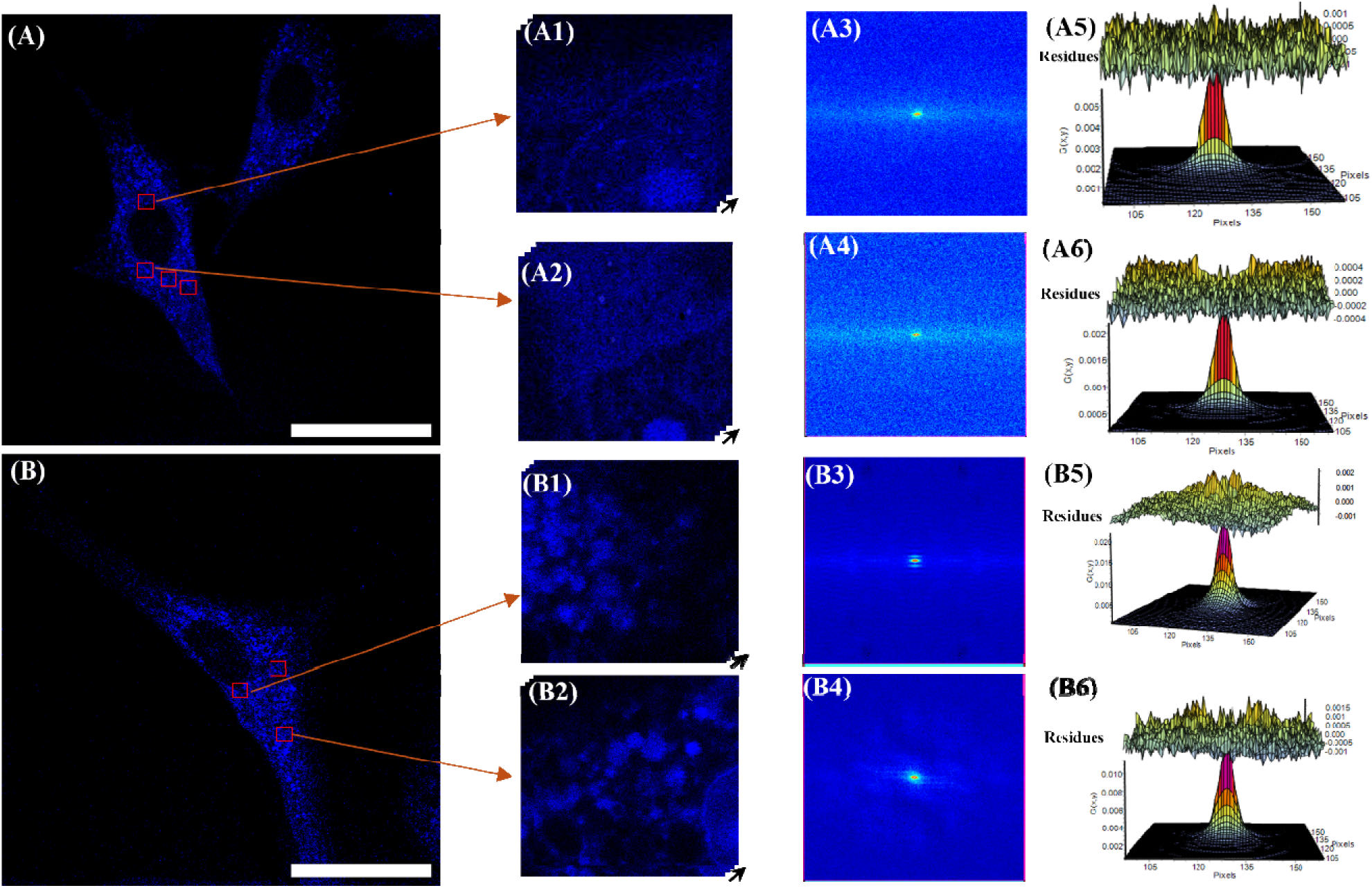
RICS on A549 cell for the calculation of diffusion measurement of CQDs and C-CQDS. (A) CQD-labeled confocal image of A549 cell (Scale bar = 40 μm). (A1) and (A2) are the magnification of the red square in A, which was used for the RICS experiment (256 x 256) at different regions of interest(ROIs). (A3) and (A4) are their corresponding 2D autocorrelation function. (A5) and (A6) are the 3D representations with residues. (B) C-CQDs labeled Confocal image of A549 cell (Scale bar = 40 μm), (B1) and (B2) are the magnification of the red square in B was used for the RICS experiment (256 x 256) at different ROIs. (B3) and (B4) are the corresponding 2D autocorrelation functions. (B5) and (B6) are the 3D representations, along with residues.

**Figure 12.**
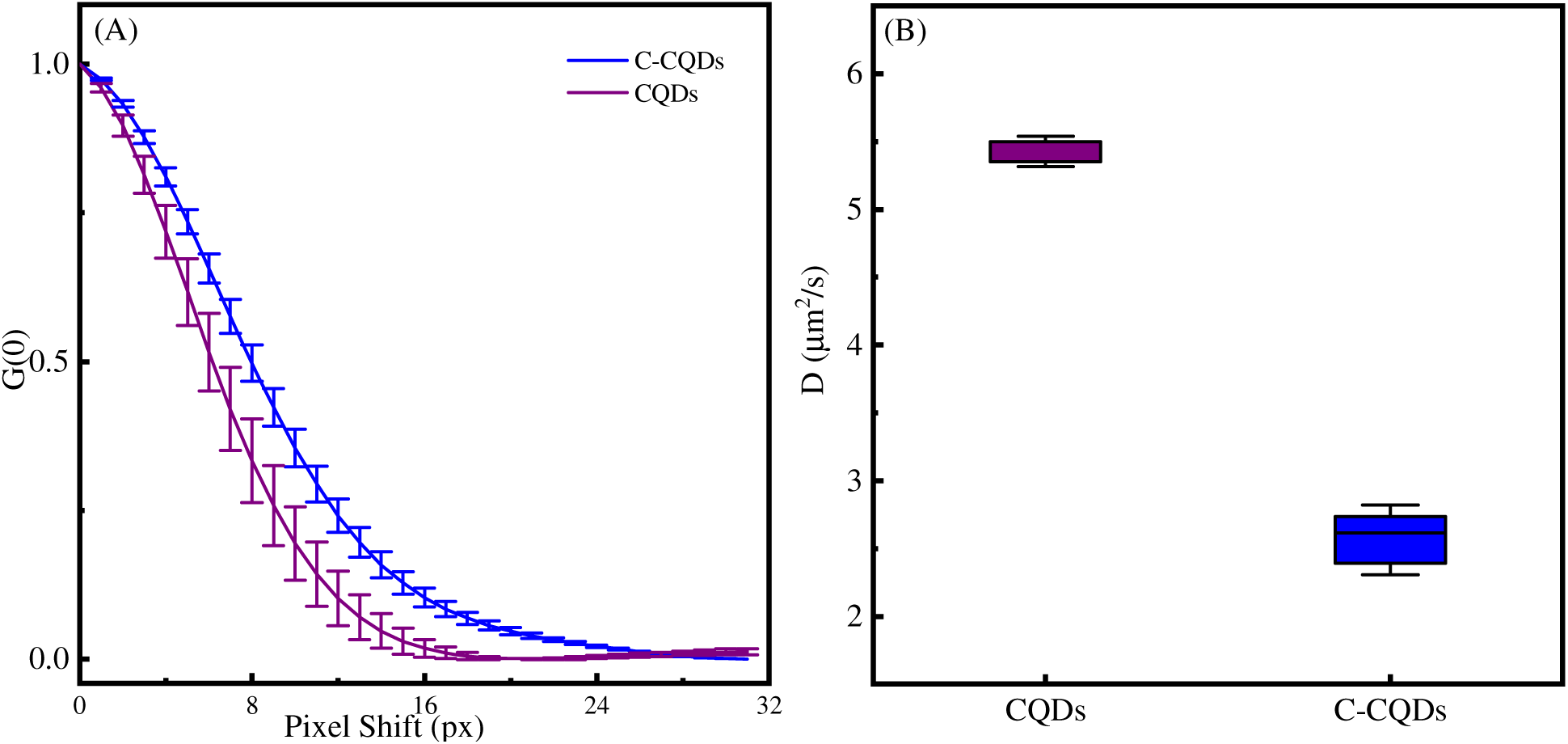
Autocorrelation function and difference in diffusion coefficient calculation. (A) Horizontal components of the autocorrelation function (ACFs) showing the difference in correlation due to CQDs and C-CQDs. Due to the smaller size of CQDs, the diffusion inside the cell is larger in CQDs, and there is a sharp decay as compared to that in C-CQDs. (B) Boxes with minimum and maximum diffusion coefficient values measured from different ROIs of a single cell, showing that C-CQDs have slower diffusion inside the cell.

The disruption of intracellular components, adhesion to cell walls, and penetration into the cells are highly modulated by the antibacterial mechanisms, and curcumin can be effective as an antibacterial agent by inducing oxidative stress, inhibiting biofilm formation, and disrupting the cell membrane.^82–83^. Here we have studied the antibacterial activity of both CQDs and C-CQDs against *E.coli* (a gram-negative bacterium) as it is considered the model organism. Upon increasing the concentration of C-CQDs from 5 to 60 μg/mL, we have observed that the growth rate of E. coli decreases gradually, whereas in the case of only CQDs, it did not show any positive results (Figure 13A). The MIC value for C-CQDs was found to be 5 μg/mL, whereas for CQDs, it is around 30 μg/mL, and the findings underscore the in-depth antibacterial properties of C-CQDs. Additionally, from the MTT assay, after 24 hours of treatment of both C-CQDs and CQDs with A549 cells, the investigation is carried out with increasing concentration of the drug. It has been found that with an increase in the concentration of C-CQDs from 5 to 40 μg/mL, it effectively decreases the % cell viability from 91% to 18% (Figure 13B). The IC_50_ value has been determined in the range of 16-17 μg/mL in the case of C-CQDs. However, after the treatment with 40 μg/mL of the drug, we have found that the effectiveness of C-CQDs is around 5 times that of CQDs.

**Figure 13.**
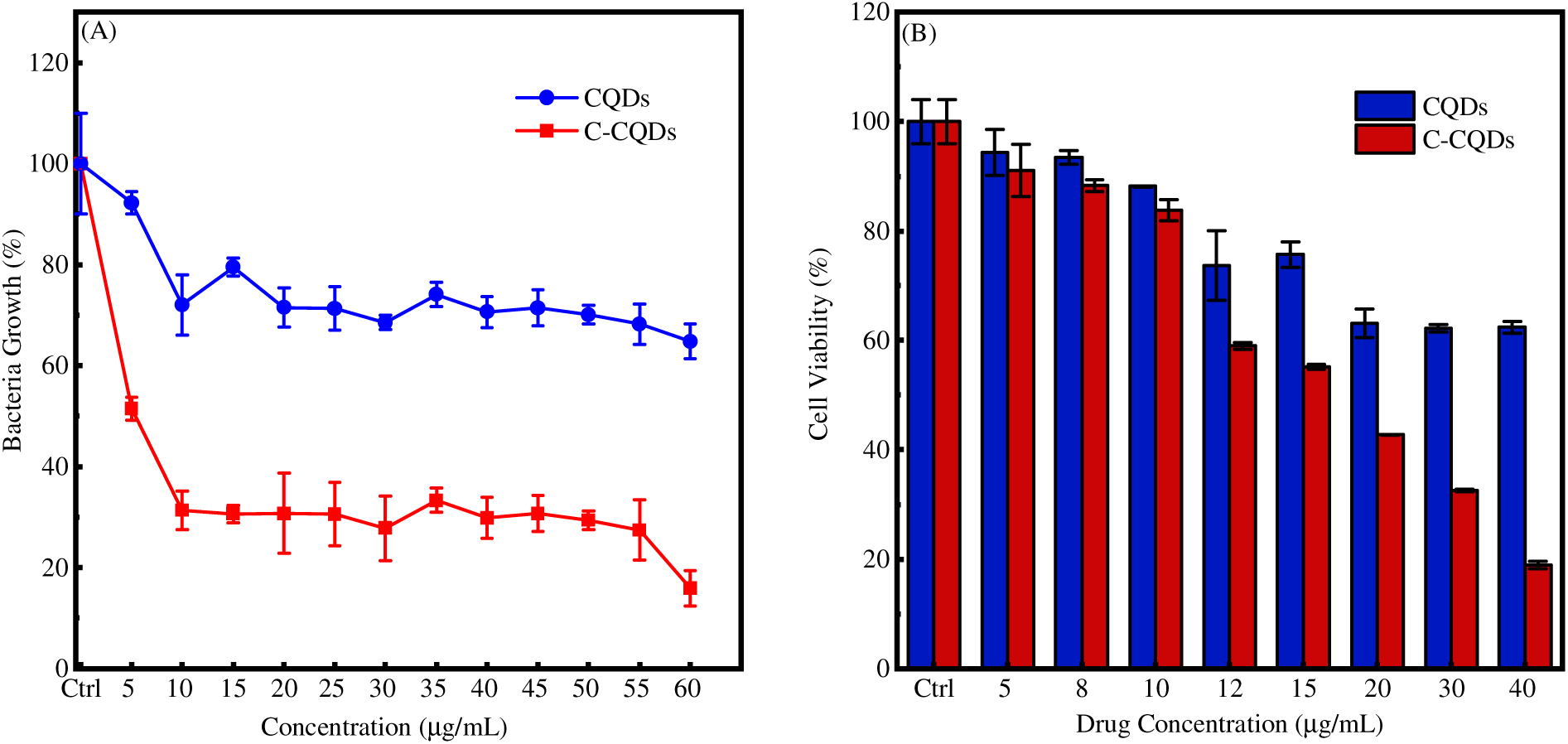
Antibacterial and cell viability study of synthesized CQDs and C-CQDs. (A) By using the MIC assay, the antibacterial study has been performed with *E. coli* bacteria (B). The MTT assay has been performed by taking A549 cells.

The appeal of the research is to know why we have used Carbon Quantum Dots and how they can conjugate the hydrophobic drug curcumin, and what type of effectiveness it can have for the cells. As compared to the commonly reported micron size of nanoparticles, the present approach is unique as the formulation of small-sized CQDs as well as C-CQDs. Due to this optimum size, it avoids lysosomal and endosomal trafficking and favors the nonendocytic delivery of the drug, unlike that of micron-sized nanoparticles, which deliver through endocytosis and phagocytosis pathways.^84–85^ As its small size has a better affinity to cross the membrane and results in cell death, so, to alleviate this issue, we can suggest using a minimum concentration as well as reducing the incubation time. Our experimental condition mimics the fluorescent nature of carbon quantum dots, broadening their applications in the field of imaging and imaging-based techniques like RICS that we have implemented in this study. As the functionalization of nanomaterials can potentially reduce their toxicity and enhance their antibacterial activity,^86^ so upon curcumin functionalization on CQDs, it shows excellent antibacterial and cytotoxicity against cells.

### Conclusions

In summary, we have demonstrated a green, eco-friendly, cost-effective, and one-step approach for synthesizing small-sized fluorescence carbon quantum dots, which can be useful for a unique drug carrier into the cytoplasm of cancerous cells. However, its excellent bioimaging probe and ability to provide a surface for the adsorption of the hydrophobic drug to enhance its solubility can be effective for targeted drug delivery. Additionally, we have employed the fluorescence techniques to investigate the interaction with proteins, which shows transient dynamic quenching and transfer of energy with bovine serum albumin protein, BSA. We have also incorporated the imaging technique, RICS, to determine the average diffusion inside the live cell for both CQDs and C-CQDs. From the RICS approach, it has been found that it follows the Stock-Einstein law, and the diffusion constant can be inversely related to the hydrodynamic diameter of the corresponding particles. Additionally, its antibacterial and cytotoxic properties are highly applicable in the field of cancer. However, this approach can be an alternative drug delivery agent and can be extended to enhance therapeutic applications.

## Supporting information

uhbnio

## Associated Content Supporting Information

The Supporting Information contains the absorption spectrum comparison, the normalised overlap spectra for FRET, and the emission spectra of BSA with CQDs only.

## Author Contributions

AKU performed all experiments, data analysis, and wrote the first draft. DKS conceived, supervised, and secured funding for the project. Both authors wrote the final version.

## ACKNOWLEDGMENT

D.K.S. thanks MoE for the STARS Grant (STARS2/2023-0473) and ANRF for the CRG Grant (ANRF/CRG/2023/003083) for the financial assistance. D.K.S. would also like to thank the financial support from the SEED Grant from IIT Jodhpur, and Start-up Research Grant from SERB (SRG/2020/001730), Govt. of India. A.K.U. would like to thank IIT Jodhpur for the fellowship. A.K.U. would also like to thank Priyanka Das, IIT Roorkee, for performing the XRD experiment and helping with the data analysis. A.K.U. would also like to thank the CRDSI facility, IIT Jodhpur, Chemistry departmental instrument facility, and AIOT-FAB facility at IIT Jodhpur for the instrumental help for the experiment.

## REFERENCES

1. Kumar, V. B.; Sher, I.; Rencus Lazar, S.; Rotenstreich, Y.; Gazit, E., Functional carbon quantum dots for ocular imaging and therapeutic applications. Small 2023, 19 (7), 2205754.

2. Wang, J.; Zhao, Y.; Sun, L.; Zhou, J.; Liu, Y.; Du, M.; Guo, Y.; Wu, X.; Li, B.; Fan, Y., Recent Advances in Carbon Dots: Applications in Oral Diseases. ACS Applied Nano Materials 2024, 8 (1), 22–38.

3. Molaei, M. J., Carbon quantum dots and their biomedical and therapeutic applications: a review. RSC advances 2019, 9 (12), 6460–6481.

4. Saini, D.; Garg, A. K.; Dalal, C.; Anand, S. R.; Sonkar, S. K.; Sonker, A. K.; Westman, G., Visible-light-promoted photocatalytic applications of carbon dots: a review. ACS Applied Nano Materials 2022, 5 (3), 3087–3109.

5. Lamba, R.; Yukta, Y.; Mondal, J.; Kumar, R.; Pani, B.; Singh, B., Carbon dots: Synthesis, characterizations, and recent advancements in biomedical, optoelectronics, sensing, and catalysis applications. ACS Applied Bio Materials 2024, 7 (4), 2086–2127.

6. Yang, Z.; Xu, T.; Li, H.; She, M.; Chen, J.; Wang, Z.; Zhang, S.; Li, J., Zero-dimensional carbon nanomaterials for fluorescent sensing and imaging. Chemical Reviews 2023, 123 (18), 11047–11136.

7. Panwar, N.; Soehartono, A. M.; Chan, K. K.; Zeng, S.; Xu, G.; Qu, J.; Coquet, P.; Yong, K.-T.; Chen, X., Nanocarbons for biology and medicine: sensing, imaging, and drug delivery. Chemical reviews 2019, 119 (16), 9559–9656.

8. Yang, L.; Wang, Z.; Wang, J.; Jiang, W.; Jiang, X.; Bai, Z.; He, Y.; Jiang, J.; Wang, D.; Yang, L., Doxorubicin conjugated functionalizable carbon dots for nucleus targeted delivery and enhanced therapeutic efficacy. Nanoscale 2016, 8 (12), 6801–6809.

9. Winer, E.; Gralow, J.; Diller, L.; Karlan, B.; Loehrer, P.; Pierce, L.; Demetri, G.; Ganz, P.; Kramer, B.; Kris, M., Clinical cancer advances 2008: major research advances in cancer treatment, prevention, and screening—a report from the American Society of Clinical Oncology. Journal of clinical oncology 2009, 27 (5), 812.

10. Ajima, K.; Murakami, T.; Mizoguchi, Y.; Tsuchida, K.; Ichihashi, T.; Iijima, S.; Yudasaka, M., Enhancement of in vivo anticancer effects of cisplatin by incorporation inside single-wall carbon nanohorns. ACS nano 2008, 2 (10), 2057–2064.

11. Quelé, L. N. d. S.; de Matos, M.; de Lima, G. G.; Brugnari, T.; Ribeiro, C. S. P.; Pedro, A. C.; Gonzalez de Cademartori, P. H.; Magalhães, W. L. E., Antimicrobial and Antioxidant Properties of Photodegraded Amorphous Curcumin on Silica Nanoparticles. ACS Applied Nano Materials 2024.

12. Banerjee, C.; Ghatak, C.; Mandal, S.; Ghosh, S.; Kuchlyan, J.; Sarkar, N., Curcumin in reverse micelle: an example to control excited-state intramolecular proton transfer (ESIPT) in confined media. The Journal of Physical Chemistry B 2013, 117 (23), 6906–6916.

13. Jovanovic, S. V.; Boone, C. W.; Steenken, S.; Trinoga, M.; Kaskey, R. B., How curcumin works preferentially with water soluble antioxidants. Journal of the American Chemical Society 2001, 123 (13), 3064–3068.

14. Kasi, P. D.; Tamilselvam, R.; Skalicka-Woźniak, K.; Nabavi, S. F.; Daglia, M.; Bishayee, A.; Pazoki-Toroudi, H.; Nabavi, S. M., Molecular targets of curcumin for cancer therapy: an updated review. Tumor Biology 2016, 37, 13017–13028.

15. Metzler, M.; Pfeiffer, E.; Schulz, S. I.; Dempe, J. S., Curcumin uptake and metabolism. Biofactors 2013, 39 (1), 14–20.

16. Hegde, M.; Girisa, S.; BharathwajChetty, B.; Vishwa, R.; Kunnumakkara, A. B., Curcumin formulations for better bioavailability: what we learned from clinical trials thus far? ACS omega 2023, 8 (12), 10713–10746.

17. Sarkar, N.; Bose, S., Liposome-encapsulated curcumin-loaded 3D printed scaffold for bone tissue engineering. ACS applied materials & interfaces 2019, 11 (19), 17184–17192.

18. Rahman, M.; Singh, J. G.; Afzal, O.; Altamimi, A. S. A.; Alrobaian, M.; Haneef, J.; Barkat, M. A.; Almalki, W. H.; Handa, M.; Shukla, R., Preparation, characterization, and evaluation of curcumin–graphene oxide complex-loaded liposomes against Staphylococcus aureus in topical disease. ACS omega 2022, 7 (48), 43499–43509.

19. Mohan Viswanathan, T.; Krishnakumar, V.; Senthilkumar, D.; Chitradevi, K.; Vijayabhaskar, R.; Rajesh Kannan, V.; Senthil Kumar, N.; Sundar, K.; Kunjiappan, S.; Babkiewicz, E., Combinatorial delivery of gallium (III) nitrate and curcumin complex-loaded hollow mesoporous silica nanoparticles for breast cancer treatment. Nanomaterials 2022, 12 (9), 1472.

20. Zhang, X.; Zhu, Y.; Fan, L.; Ling, J.; Yang, L.-Y.; Wang, N.; Ouyang, X.-k., Delivery of curcumin by fucoidan-coated mesoporous silica nanoparticles: Fabrication, characterization, and in vitro release performance. International Journal of Biological Macromolecules 2022, 211, 368–379.

21. Wei, T.; Zhang, Y.; Lei, M.; Qin, Y.; Wang, Z.; Chen, Z.; Zhang, L.; Zhu, Y., Development of oral curcumin based on pH-responsive transmembrane peptide-cyclodextrin derivative nanoparticles for hepatoma. Carbohydrate Polymers 2022, 277, 118892.

22. Kim, C. H.; Kim, B. D.; Lee, T. H.; Kim, H. K.; Lyu, M. J.; Yoon, Y. I.; Goo, Y. T.; Kang, M. J.; Lee, S.; Choi, Y. W., Synergistic co-administration of docetaxel and curcumin to chemoresistant cancer cells using PEGylated and RIPL peptide-conjugated nanostructured lipid carriers. Cancer Nanotechnology 2022, 13 (1), 17.

23. Souto, E. B.; Ribeiro, A. F.; Ferreira, M. I.; Teixeira, M. C.; Shimojo, A. A.; Soriano, J. L.; Naveros, B. C.; Durazzo, A.; Lucarini, M.; Souto, S. B., New nanotechnologies for the treatment and repair of skin burns infections. International journal of molecular sciences 2020, 21 (2), 393.

24. Le, N.; Zhang, M.; Kim, K., Quantum dots and their interaction with biological systems. International Journal of Molecular Sciences 2022, 23 (18), 10763.

25. Zhang, B.; Wang, X.; Liu, F.; Cheng, Y.; Shi, D., Effective reduction of nonspecific binding by surface engineering of quantum dots with bovine serum albumin for cell-targeted imaging. Langmuir 2012, 28 (48), 16605–16613.

26. Lu, R.; Li, W.-W.; Katzir, A.; Raichlin, Y.; Yu, H.-Q.; Mizaikoff, B., Probing the secondary structure of bovine serum albumin during heat-induced denaturation using mid-infrared fiberoptic sensors. Analyst 2015, 140 (3), 765–770.

27. Patel, A.; Paul, S.; Akhtar, N.; Das, S.; Kar, S.; Bhattacharjee, S.; Manna, D., Onium-and alkyl amine-decorated protein nanoparticles as antimicrobial agents and carriers of antibiotics to promote synergistic antibacterial and antibiofilm activities. ACS Applied Nano Materials 2022, 5 (11), 16602–16611.

28. Ahmad, M.; Singla, N.; Bhadwal, S. S.; Kaur, S.; Singh, P.; Kumar, S., Differentiation of HSA and BSA and Instantaneous Detection of HSO3–Using Confined Space of Serum Albumins and Live Cell Imaging of Exogenous/Endogenous HSO3–. ACS omega 2023, 8 (2), 2639–2647.

29. Dennison, J. M.; Zupancic, J. M.; Lin, W.; Dwyer, J. H.; Murphy, C. J., Protein adsorption to charged gold nanospheres as a function of protein deformability. Langmuir 2017, 33 (31), 7751–7761.

30. Yin, M.-M.; Dong, P.; Chen, W.-Q.; Xu, S.-P.; Yang, L.-Y.; Jiang, F.-L.; Liu, Y., Thermodynamics and mechanisms of the interactions between ultrasmall fluorescent gold nanoclusters and human serum albumin, γ-globulins, and transferrin: a spectroscopic approach. Langmuir 2017, 33 (21), 5108–5116.

31. Agarwala, P.; Bera, T.; Sasmal, D. K., Molecular Mechanism of Interaction of Curcumin with BSA, Surfactants and Live E. Coli Cell Membrane Revealed by Fluorescence Spectroscopy and Confocal Microscopy. ChemPhysChem 2022, 23 (18), e202200265.

32. Digman, M. A.; Gratton, E., Analysis of diffusion and binding in cells using the RICS approach. Microscopy research and technique 2009, 72 (4), 323–332.

33. Rossow, M. J.; Sasaki, J. M.; Digman, M. A.; Gratton, E., Raster image correlation spectroscopy in live cells. Nature protocols 2010, 5 (11), 1761–1774.

34. Gielen, E.; Smisdom, N.; vandeven, M.; De Clercq, B.; Gratton, E.; Digman, M.; Rigo, J.-M.; Hofkens, J.; Engelborghs, Y.; Ameloot, M., Measuring diffusion of lipid-like probes in artificial and natural membranes by raster image correlation spectroscopy (RICS): use of a commercial laser-scanning microscope with analog detection. Langmuir 2009, 25 (9), 5209–5218.

35. Brown, C.; Dalal, R.; Hebert, B.; Digman, M.; Horwitz, A.; Gratton, E., Raster image correlation spectroscopy (RICS) for measuring fast protein dynamics and concentrations with a commercial laser scanning confocal microscope. Journal of microscopy 2008, 229 (1), 78–91.

36. Rout, D.; Upadhyaya, A. K.; Agarwala, P.; Sharma, C.; Pal, A.; Sasmal, D. K., Drug Binding to Partially Unfolded Serum Albumin: Insights into Nonsteroidal Anti-Inflammatory Drug Naproxen–BSA Interactions from Spectroscopic and MD Simulation Studies. The Journal of Physical Chemistry B 2024, 128 (39), 9327–9340.

37. Sharma, S.; Banjare, M. K.; Singh, N.; Korábečný, J.; Kuča, K.; Ghosh, K. K., Multi-spectroscopic monitoring of molecular interactions between an amino acid-functionalized ionic liquid and potential anti-Alzheimer’s drugs. RSC advances 2020, 10 (64), 38873–38883.

38. Upadhyaya, A. K.; Agarwala, P.; Sharma, C.; Sasmal, D. K., Synthesis and Characterization of N Doped Carbon Quantum Dots and its Application for Efficient Delivery of Curcumin in Live Cell. ChemPhysChem 2025, 26 (6), e202400855.

39. Magde, D.; Elson, E. L.; Webb, W. W., Fluorescence correlation spectroscopy. II. An experimental realization. Biopolymers: Original Research on Biomolecules 1974, 13 (1), 29–61.

40. Mohammad, M.; Saha, I.; Pal, K.; Karmakar, P.; Pandya, P.; Gazi, H. A. R.; Islam, M. M., A comparison on the biochemical activities of Fluorescein disodium, Rose Bengal and Rhodamine 101 in the light of DNA binding, antimicrobial and cytotoxic study. Journal of Biomolecular Structure and Dynamics 2022, 40 (20), 9848–9859.

41. Neufeld, B. H.; Tapia, J. B.; Lutzke, A.; Reynolds, M. M., Small molecule interferences in resazurin and MTT-based metabolic assays in the absence of cells. Analytical chemistry 2018, 90 (11), 6867–6876.

42. Kubin, R. F.; Fletcher, A. N., Fluorescence quantum yields of some rhodamine dyes. Journal of Luminescence 1982, 27 (4), 455–462.

43. Rout, D.; Sharma, S.; Agarwala, P.; Upadhyaya, A. K.; Sharma, A.; Sasmal, D. K., Interaction of Ibuprofen with Partially Unfolded Bovine Serum Albumin in the Presence of Ionic Micelles and Oligosaccharides at Different λex and pH: A Spectroscopic Analysis. ACS omega 2023, 8 (3), 3114–3128.

44. Boruah, J. S.; Sankaranarayanan, K.; Chowdhury, D., Insight into carbon quantum dot– vesicles interactions: role of functional groups. RSC advances 2022, 12 (7), 4382–4394.

45. Tyagi, A.; Tripathi, K. M.; Singh, N.; Choudhary, S.; Gupta, R. K., Green synthesis of carbon quantum dots from lemon peel waste: applications in sensing and photocatalysis. RSC advances 2016, 6 (76), 72423–72432.

46. Kumar, A.; Kumar, I.; Gathania, A. K., Synthesis, characterization and potential sensing application of carbon dots synthesized via the hydrothermal treatment of cow milk. Scientific Reports 2022, 12 (1), 22495.

47. Cheng, Y.; Qiang, S.; Li, J.; Wei, W.; Kuang, Y.; Zhang, W.; Fang, X.; Ding, T.; Guo, L.; Chen, Y., Assembly of Centimeter-Scale Plasmonic Nanocavities for Bright and Ultrafast Emission of Red Carbon Dots. ACS Applied Nano Materials 2022, 5 (10), 14902–14911.

48. Yang, C.-R.; Tseng, S.-F.; Chen, Y.-T., Characteristics of graphene oxide films reduced by using an atmospheric plasma system. Nanomaterials 2018, 8 (10), 802.

49. Tremblay, C.; Booth, I.; Thomas, S.; Cordoba, C.; Blackburn, A. M.; Buckley, H. L., Application of Hydrogenated Graphitic Supports in Electrocatalysts: Effects on Carbon Support Surface Chemistry, Nanoparticle Growth, and Electrocatalytic Activity. ACS Applied Materials & Interfaces 2025.

50. Guo, H.; Zhang, J.; Gao, J.; Liang, Z.; Li, W.; Yan, H.; Guo, R.; Wang, H., Carbon quantum dots-modified TiO2 nanoparticles for antibacterial applications. Chemical Physics Letters 2025, 869, 142055.

51. Lotey, N. K. B.; Lemos, R.; D’Silva, F.; Deshmukh, A.; Singh, N.; Wankhede, S.; Davuluri, R.; Vishe, N.; Kulkarni, S., Highly Fluorescent Bio-Synthesized Carbon Quantum Dots for Latent Fingerprint Detection. Journal of Fluorescence 2025, 1–11.

52. Wu, P.; Li, W.; Wu, Q.; Liu, Y.; Liu, S., Hydrothermal synthesis of nitrogen-doped carbon quantum dots from microcrystalline cellulose for the detection of Fe 3+ ions in an acidic environment. RSC advances 2017, 7 (70), 44144–44153.

53. Ossonon, B. D.; Bélanger, D., Synthesis and characterization of sulfophenyl-functionalized reduced graphene oxide sheets. RSC advances 2017, 7 (44), 27224–27234.

54. Saini, S.; Saini, P.; Kumar, K.; Sethi, M.; Meena, P.; Gurjar, A.; Dandia, A.; Dhuria, T.; Parewa, V., Unlocking the molecular behavior of natural amine-targeted carbon quantum dots for the synthesis of diverse pharmacophore scaffolds via an unusual nanoaminocatalytic route. ACS Applied Materials & Interfaces 2023, 15 (42), 49083–49094.

55. Mondal, T. K.; Kapuria, A.; Miah, M.; Saha, S. K., Solubility tuning of alkyl amine functionalized carbon quantum dots for selective detection of nitroexplosive. Carbon 2023, 209, 117972.

56. Mecozzi, M.; Pietroletti, M.; Scarpiniti, M.; Acquistucci, R.; Conti, M. E., Monitoring of marine mucilage formation in Italian seas investigated by infrared spectroscopy and independent component analysis. Environmental Monitoring and Assessment 2012, 184, 6025–6036.

57. Ferjani, H.; Abdalla, S.; Oyewo, O. A.; Onwudiwe, D. C., Facile synthesis of carbon dots by the hydrothermal carbonization of avocado peels and evaluation of the photocatalytic property. Inorganic Chemistry Communications 2024, 160, 111866.

58. Edison, T. N. J. I.; Atchudan, R.; Sethuraman, M. G.; Shim, J.-J.; Lee, Y. R., Microwave assisted green synthesis of fluorescent N-doped carbon dots: Cytotoxicity and bio-imaging applications. Journal of Photochemistry and Photobiology B: Biology 2016, 161, 154–161.

59. Korin, E.; Froumin, N.; Cohen, S., Surface analysis of nanocomplexes by X-ray photoelectron spectroscopy (XPS). ACS Biomaterials Science & Engineering 2017, 3 (6), 882–889.

60. Elkun, S.; Ghali, M.; Sharshar, T.; Mosaad, M. M., Green synthesis of fluorescent N-doped carbon quantum dots from castor seeds and their applications in cell imaging and pH sensing. Scientific Reports 2024, 14 (1), 27927.

61. Dager, A.; Uchida, T.; Maekawa, T.; Tachibana, M., Synthesis and characterization of mono-disperse carbon quantum dots from fennel seeds: photoluminescence analysis using machine learning. Scientific reports 2019, 9 (1), 14004.

62. Nguyen, K. G.; Baragau, I.-A.; Gromicova, R.; Nicolaev, A.; Thomson, S. A.; Rennie, A.; Power, N. P.; Sajjad, M. T.; Kellici, S., Investigating the effect of N-doping on carbon quantum dots structure, optical properties and metal ion screening. Scientific Reports 2022, 12 (1), 13806.

63. Baliyan, A.; Nakajima, Y.; Fukuda, T.; Uchida, T.; Hanajiri, T.; Maekawa, T., Synthesis of an ultradense forest of vertically aligned triple-walled carbon nanotubes of uniform diameter and length using hollow catalytic nanoparticles. Journal of the American Chemical Society 2014, 136 (3), 1047–1053.

64. Chellasamy, G.; Ankireddy, S. R.; Lee, K.-N.; Govindaraju, S.; Yun, K., Smartphone-integrated colorimetric sensor array-based reader system and fluorometric detection of dopamine in male and female geriatric plasma by bluish-green fluorescent carbon quantum dots. Materials Today Bio 2021, 12, 100168.

65. Wang, S.; Li, L.; Zhu, Z.; Zhao, M.; Zhang, L.; Zhang, N.; Wu, Q.; Wang, X.; Li, G., Remarkable improvement in photocatalytic performance for tannery wastewater processing via SnS2 modified with N doped carbon quantum dots: synthesis, characterization, and 4 nitrophenol aided Cr (VI) photoreduction. Small 2019, 15 (29), 1804515.

66. Mishra, J.; Suryawanshi, T.; Redkar, N.; Kumar Das, R.; Saxena, S.; Majumder, A.; Kondabagil, K.; Shukla, S., Toxicological Effects of Metal Doped Carbon Quantum Dots. ChemSusChem 2025, e202402056.

67. Alqahtani, A.; Alqahtani, T.; Al Fatease, A.; Alshehri, A.; Almrasy, A. A., Development of N, P-doped carbon quantum dots as a green fluorescent probe for fexofenadine determination: mechanistic studies, Box–Behnken optimization, and pharmacokinetic application. RSC advances 2025, 15 (18), 14545–14557.

68. Kayyal, T. B.; Tucker, J.; Lowrance, C. M.; Ajiboye, L.; Pelton, M.; Bennett, J. W.; Daniel, M.-C., Oleic acid rearrangement enables facile transfer of red-emitting quantum dots from hexane into water with enhanced fluorescence. Nanoscale 2025.

69. Zaini, M. S.; Liew, J. Y. C.; Paiman, S.; Tee, T. S.; Kamarudin, M. A., Solvent-dependent photoluminescence emission and colloidal stability of carbon quantum dots from watermelon peels. Journal of Fluorescence 2025, 35 (1), 245–256.

70. Dager, A.; Baliyan, A.; Kurosu, S.; Maekawa, T.; Tachibana, M., Ultrafast synthesis of carbon quantum dots from fenugreek seeds using microwave plasma enhanced decomposition: application of C-QDs to grow fluorescent protein crystals. Scientific reports 2020, 10 (1), 12333.

71. Setianto, S.; Men, L. K.; Bahtiar, A.; Panatarani, C.; Joni, I. M., Carbon quantum dots with honeycomb structure: a novel synthesis approach utilizing cigarette smoke precursors. Scientific Reports 2024, 14 (1), 1996.

72. Singh, A. K.; Prakash, P.; Singh, R.; Nandy, N.; Firdaus, Z.; Bansal, M.; Singh, R. K.; Srivastava, A.; Roy, J. K.; Mishra, B., Curcumin quantum dots mediated degradation of bacterial biofilms. Frontiers in microbiology 2017, 8, 1517.

73. Su, R.; Yan, H.; Jiang, X.; Zhang, Y.; Li, P.; Su, W., Orange-red to NIR emissive carbon dots for antimicrobial, bioimaging and bacteria diagnosis. Journal of materials chemistry B 2022, 10 (8), 1250–1264.

74. Anitha, A.; Deepagan, V.; Rani, V. D.; Menon, D.; Nair, S.; Jayakumar, R., Preparation, characterization, in vitro drug release and biological studies of curcumin loaded dextran sulphate–chitosan nanoparticles. Carbohydrate Polymers 2011, 84 (3), 1158–1164.

75. Yallapu, M. M.; Jaggi, M.; Chauhan, S. C., β-Cyclodextrin-curcumin self-assembly enhances curcumin delivery in prostate cancer cells. Colloids and surfaces B: Biointerfaces 2010, 79 (1), 113–125.

76. Nguyen, T. A.; Tang, Q. D.; Doan, D. C. T.; Dang, M. C., Micro and nano liposome vesicles containing curcumin for a drug delivery system. Advances in Natural Sciences: Nanoscience and Nanotechnology 2016, 7 (3), 035003.

77. Khatun, M. A.; Sultana, F.; Saha, I.; Karmakar, P.; Gazi, H. A. R.; Islam, M. M.; Show, B.; Mukhopadhyay, S., Lentil Extract-Mediated Ag QD Synthesis: Predilection for Albumin Protein Interaction, Antibacterial Activity, and Its Cytotoxicity for Wi-38 and PC-3 Cell Lines. ACS Applied Bio Materials 2024, 7 (10), 6568–6582.

78. Ghosh, G.; Panicker, L., Protein–nanoparticle interactions and a new insight. Soft Matter 2021, 17 (14), 3855–3875.

79. Boehmler, D. J.; O’Dell, Z. J.; Chung, C.; Riley, K. R., Bovine serum albumin enhances silver nanoparticle dissolution kinetics in a size-and concentration-dependent manner. Langmuir 2020, 36 (4), 1053–1061.

80. Sahoo, R.; Mukherjee, N.; Paramanik, S.; Jana, N. R., Vegetable Oil–Based Pickering Nanoemulsions As Carriers for Cytosolic Drug Delivery. ACS Applied Nano Materials 2024, 7 (13), 15702–15709.

81. Seemork, J.; Sansureerungsikul, T.; Sathornsantikun, K.; Sinthusake, T.; Shigyou, K.; Tree-Udom, T.; Jiangchareon, B.; Chiablaem, K.; Lirdprapamongkol, K.; Svasti, J., Penetration of oxidized carbon nanospheres through lipid bilayer membrane: comparison to graphene oxide and oxidized carbon nanotubes, and effects of pH and membrane composition. ACS applied materials & interfaces 2016, 8 (36), 23549–23557.

82. Hussain, Y.; Alam, W.; Ullah, H.; Dacrema, M.; Daglia, M.; Khan, H.; Arciola, C. R., Antimicrobial potential of curcumin: therapeutic potential and challenges to clinical applications. Antibiotics 2022, 11 (3), 322.

83. Dai, C.; Lin, J.; Li, H.; Shen, Z.; Wang, Y.; Velkov, T.; Shen, J., The natural product curcumin as an antibacterial agent: Current achievements and problems. Antioxidants 2022, 11 (3), 459.

84. Xia, Y.; Wu, J.; Wei, W.; Du, Y.; Wan, T.; Ma, X.; An, W.; Guo, A.; Miao, C.; Yue, H., Exploiting the pliability and lateral mobility of Pickering emulsion for enhanced vaccination. Nature materials 2018, 17 (2), 187–194.

85. Du, Y.; Song, T.; Wu, J.; Gao, X.-D.; Ma, G.; Liu, Y.; Xia, Y., Engineering mannosylated pickering emulsions for the targeted delivery of multicomponent vaccines. Biomaterials 2022, 280, 121313.

86. Barbalinardo, M.; Bertacchini, J.; Bergamini, L.; Magarò, M. S.; Ortolani, L.; Sanson, A.; Palumbo, C.; Cavallini, M.; Gentili, D., Surface properties modulate protein corona formation and determine cellular uptake and cytotoxicity of silver nanoparticles. Nanoscale 2021, 13 (33), 14119–14129.

