## Supplementary material for "Curcumin Conjugated Carbon Quantum Dots: A Theranostic Probe to Study BSA Interaction, Cellular Imaging, and RICS-Based Intracellular Transport": uhbnio

Jodhpur, Rajasthan 343037, India

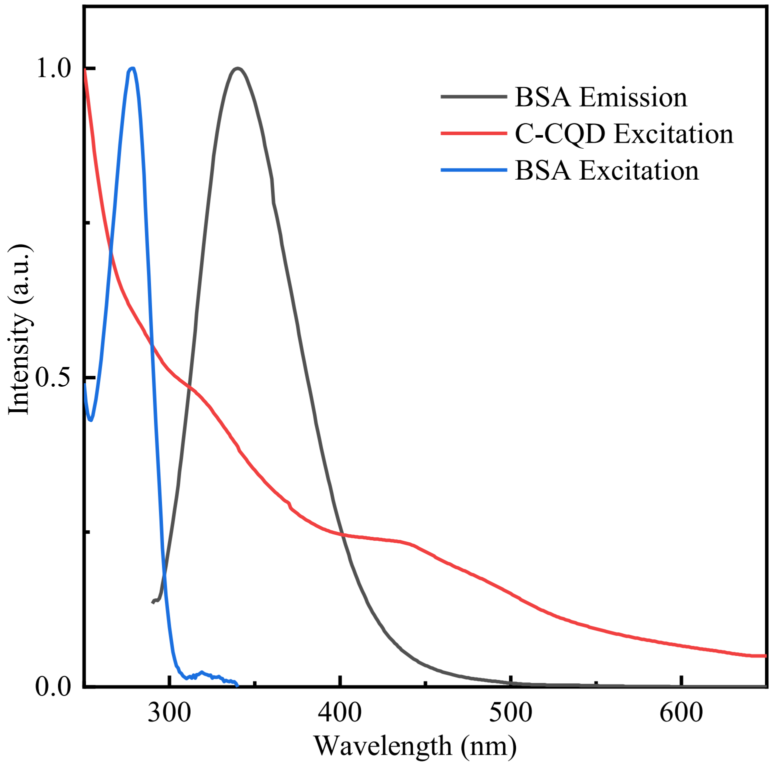


**Figure S1. The normalized Overlap excitation and emission spectra of Donors and acceptors for the FRET study.** From the FRET images, it is indicated that the emission of donor (BSA) overlaps with the excitation spectra of acceptor (C-CQDs)

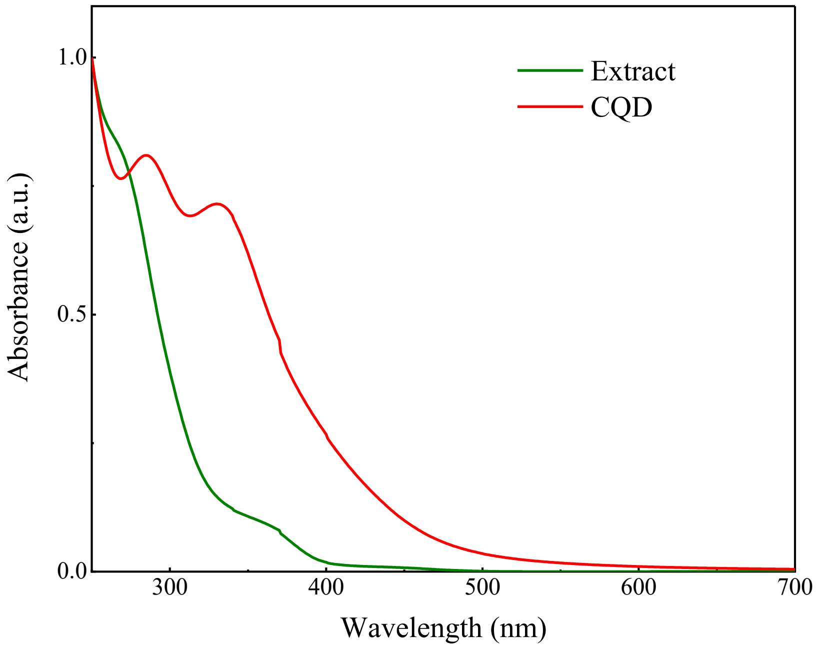


**Figure S2. The normalized excitation spectra of CQDs in comparison with only the leaf extract.** It has been found that there is no such defined peak in the case of only the extract. But in the case of CQDs, two well-defined peaks are there because of n-p$^*$ and p-p$^*$ transitions.

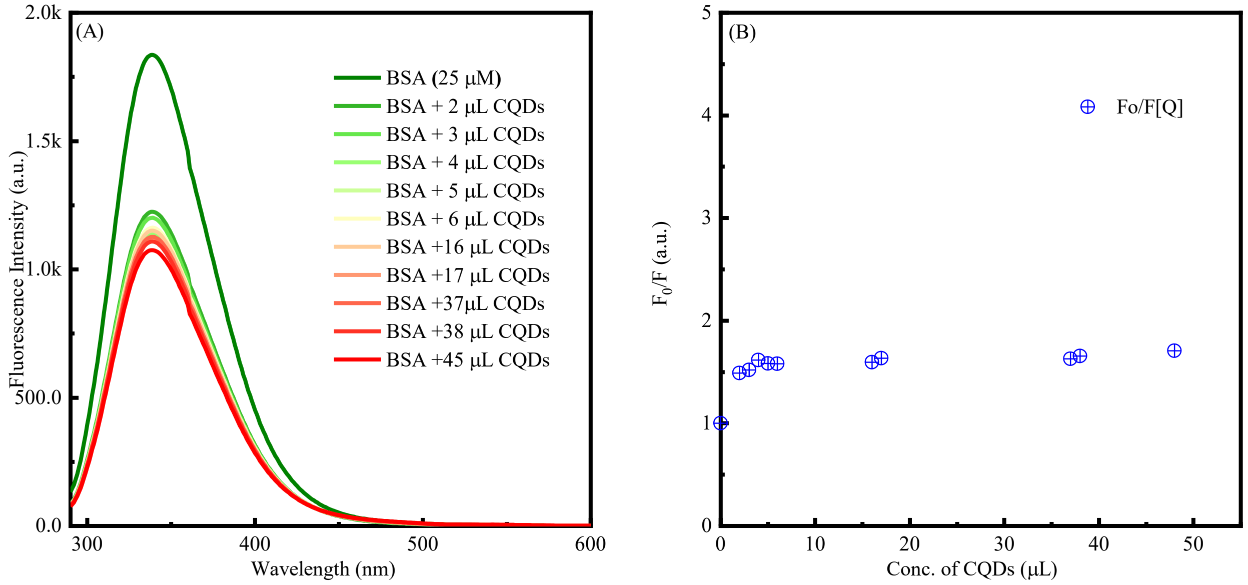


**Figure S3. Emission spectra of BSA with CQDs.** (A)The results show that there is no quenching of fluorescence upon the addition of only CQDs. However, there is a sharp decrease in fluorescence with the addition of 2 mL of CQDs, which may be due to the effect of CQDs on BSA. (B) The corresponding Benesi-Hildebrand plots show no linear relationship.
